# Structural insights into GrpEL1-mediated nucleotide and substrate release of human mitochondrial Hsp70

**DOI:** 10.1101/2024.05.10.593630

**Authors:** Marc A. Morizono, Kelly L. McGuire, Natalie I. Birouty, Mark A. Herzik

## Abstract

Maintenance of protein homeostasis is necessary for cell viability and depends on a complex network of chaperones and co-chaperones, including the heat-shock protein 70 (Hsp70) system. In human mitochondria, mitochondrial Hsp70 (mortalin) and the nucleotide exchange factor (GrpEL1) work synergistically to stabilize proteins, assemble protein complexes, and facilitate protein import. However, our understanding of the molecular mechanisms guiding these processes is hampered by limited structural information. To elucidate these mechanistic details, we used cryoEM to determine the first structures of full-length human mortalin-GrpEL1 complexes in previously unobserved states. Our structures and molecular dynamics simulations allow us to delineate specific roles for mortalin-GrpEL1 interfaces and to identify steps in GrpEL1-mediated nucleotide and substrate release by mortalin. Subsequent analyses reveal conserved mechanisms across bacteria and mammals and facilitate a complete understanding of sequential nucleotide and substrate release for the Hsp70 chaperone system.

## Introduction

Cells maintain the functional states, relative abundance, and proper localization of their myriad proteins through a complex regulatory network governing all aspects of protein synthesis, folding, quality control, and degradation that is collectively termed proteostasis.^1,2^ Molecular chaperone systems like heat-shock protein 70 (Hsp70) and Hsp90 proteins play key roles in proteostasis by mediating the initial folding or refolding of many proteins, regulating their biochemical activities, and guiding them to their appropriate destinations in the cell.^3,4^ Typically, molecular chaperone systems are characterized by the binding of exposed hydrophobic regions and are commonly coupled with an array of co-chaperones that aid in the regulation of their activity and specificity.^3–5^ Members of the Hsp70 family are among the most ubiquitous molecular chaperones and are found in all kingdoms of life.^6,7^ Hsp70s exhibit promiscuous binding of short hydrophobic sequences within client proteins and use ATPase activity to ensure that protein folding pathways, translocation events, protein degradation, and signaling cascades are maintained in both basal and stressed cellular environments.^3–6,8^ Unsurprisingly, several diseases and cancers have been attributed to the dysfunction of Hsp70 activity.^9–12^ Hsp70s also collaborate with two classes of co-chaperones, termed J-proteins (Hsp40s) and nucleotide exchange factors (NEFs), that stimulate ATP hydrolysis and release of ADP/substrate, respectively **(Figure 1A)**.^13–16^ These co-chaperones are critical for Hsp70 function as they endow substrate specificity and catalytic efficiency to an otherwise promiscuous and inefficient ATPase.^5,13–16^ In the canonical Hsp70 substrate capture and release cycle, Hsp70 binds ATP at its N-terminal nucleotide-binding domain (NBD) and adopts a compacted conformation, with the C-terminal substrate binding domain (SBD) interacting substantially with the NBD **(Figure 1A**,**B)**. Upon binding of a protein substrate at the SBD, Hsp70 cooperates with a stimulatory J-protein to hydrolyze ATP and release inorganic phosphate, adopting an extended conformation with the ADP-bound NBD and substrate-bound SBD separated via the conserved interdomain linker (IDL).^16–19^ In bacteria, mitochondria, and chloroplasts, the GrpE-family of NEFs interact with substrate-bound Hsp70 to facilitate ADP and protein substrate release.^16,20–23^ ATP binding results in rapid dissociation of the NEF and enables release of nucleotide and substrate, thus resetting Hsp70 to its ATP-bound state **(Figure 1A)**.^20,21,24^ In mitochondria and chloroplasts, Hsp70s and their co-chaperones mediate not only protein folding and protein complex assembly but also are essential for translocation of proteins across the membranes of these organelles.^16,19,25,26^ Mammalian mitochondria harbor a distinctive Hsp70 homolog (mtHsp70) known as mortalin (*Hs*HSPA9 in humans).^27–29^ Mortalin and the human mitochondrial NEF, GrpEL1, together facilitate several key biological processes within the mitochondrial matrix **(Figure 1B)**.^30–33^ Mortalin and GrpEL1 work with two distinct J-protein complexes to perform their functions. The DNAJA3 complex mediates protein folding and protein complex assembly,^34–36^ while the Pam16/Pam18 complex aids translocation of proteins through the translocase of the inner mitochondrial membrane-23 (TIM23) complex, via the presequence associated motor (PAM) complex.^16,37–39^

**Figure 1:**
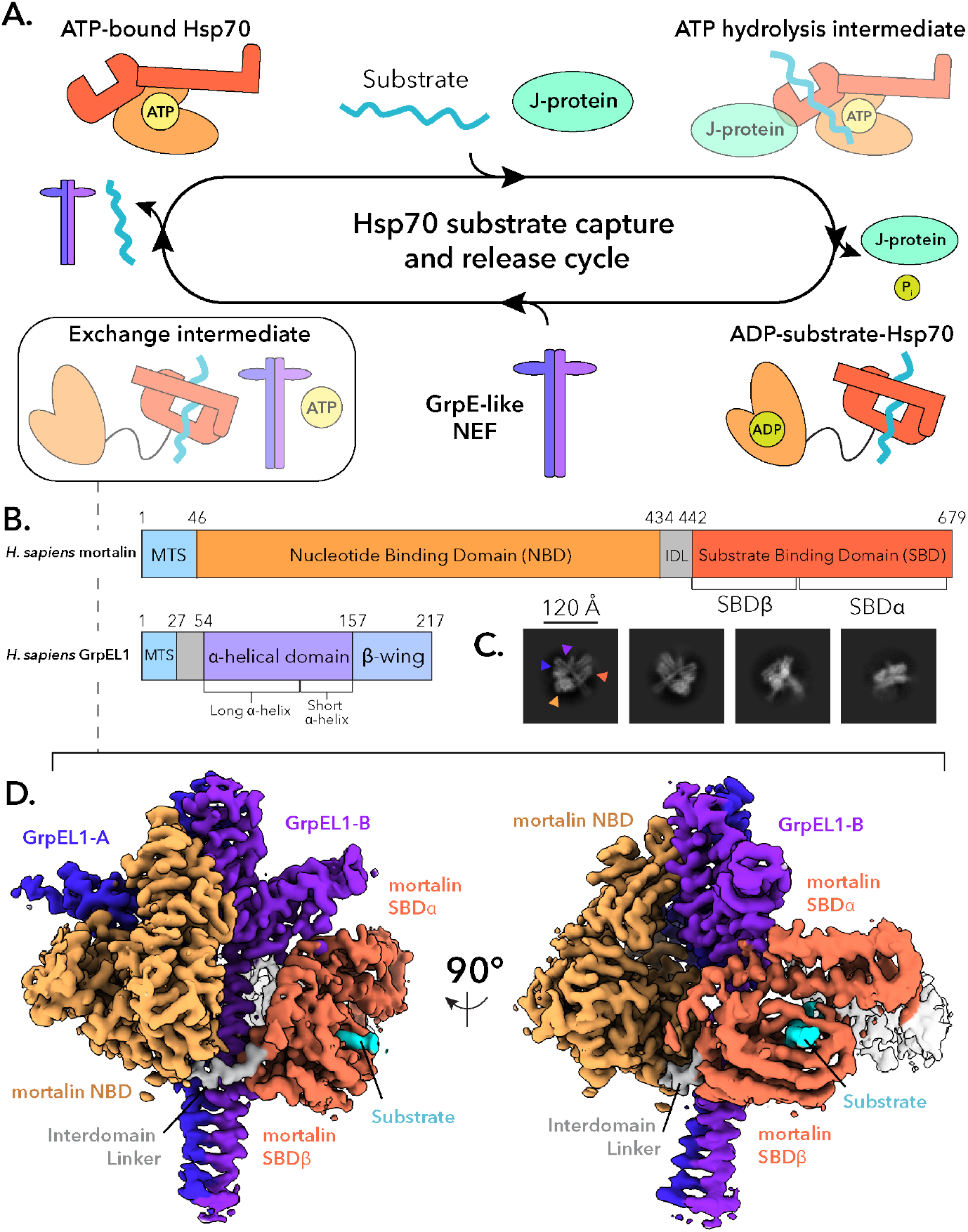
Structural determination of the *Hs*mortalin-GrpEL1 complex. **A**. Canonical Hsp70 substrate capture and release cycle. **B**. Domain topology of *Hs*mortalin and *Hs*GrpEL1. MTS: mitochondrial targeting signal. IDL: interdomain linker. **C**. Representative D classes of the *Hs*mortalin-GrpEL1 complex. Colored arrows correspond to the densities observed in **D. D**. DeepEMhancer-sharpened map of the *Hs*mortalin-GrpEL1 complex colored by subunits and subdomains. The structure represents the exchange intermediate depicted in **A**.

Insights into mortalin–GrpEL1 function have come primarily from studies of their bacterial counterparts, DnaK and GrpE.^21,24,40,41^ While these studies have revealed a putative mechanism for GrpE-mediated nucleotide release, existing structures of bacterial proteins are limited by use of truncated constructs or flexible and disordered regions.^22,40,42^ These limitations have not only precluded our understanding of critical substrate binding and release allosteric mechanisms, but also hinder elucidation of these mechanisms in eukaryotic contexts. Thus, comprehensive structural and mechanistic studies of eukaryotic Hsp70s, in particular mortalin with its co-chaperones, are needed.

To elucidate the mechanisms guiding nucleotide and substrate exchange in mortalin, we performed cryoEM structural analysis of human mortalin–GrpEL1 complexes. Herein, we present the first visualization of full-length human mortalin in a nucleotide-free and substrate-bound state in complex with full-length GrpEL1. Comparisons with bacterial DnaK and GrpE structures reveal new insights into the roles of GrpE-family NEFs in mediating nucleotide and substrate release. Finally, we combine cryoEM and molecular dynamics (MD) to characterize an interface between GrpEL1 and the mortalin SBD that likely plays a key role in substrate release. Overall, this work enables comparative analyses between bacterial DnaK-GrpE and human mortalin-GrpEL1 chaperone systems and points to a conserved mechanism for GrpE-family NEFs in nucleotide and substrate release from Hsp70-family chaperones.

## Results

### Structure of the human mortalin-GrpEL1 complex

To better understand how eukaryotic Hsp70 proteins dynamically interact with their co-chaperones and substrates, we reconstituted a complex of full-length human mortalin and GrpEL1 with a soluble chimeric construct of ^2-^ Pam16/Pam18 (see ***Methods*** and **Supplementary Figure 1**). To stabilize the complex for structural analysis, we introduced a point mutation, R126W, into mortalin (mortalin_*R*126*W*_, hereafter referred to as mortalin) that is associated with EVEN-PLUS syndrome and has been shown to reduce mortalin’s ATPase activity and conformational flexibility.^12^ Attempts to form a mortalin–GrpEL1 complex with wild-type mortalin resulted in inconsistent complex formation, whereas mixing GrpEL1 with mortalin_*R*126*W*_ resulted in reproducible complex. Incubation of these components and subsequent separation via size exclusion chromatography yielded a high molecular weight species that appeared to comprise mortalin, GrpEL1, and a lower molecular weight species **(Supplementary Figure 2)**. While Pam16/Pam18 is not bound to this complex, its inclusion in the protein mixture was nonetheless important to mediate proper assembly of the stable mortalin-GrpEL1 complex. We performed cryoEM analysis of the mortalin–GrpEL1 complex which yielded two-dimensional (2-D) class averages representing the canonical GrpE-like extended dimer with apparent density for mortalin on both sides of the *α*-helical stalk domain **(Figure 1C)**. Initial reconstructions yielded a cryoEM density containing the mortalin NBD and GrpEL1 dimer, however, density for the substrate binding domain (SBD) was weak. Heterogeneous refinement using reference volumes with and without SBD density allowed separation of particles that contained the entirety of the SBD **(Supplementary Figure 3)**. Further refinement yielded a 2.96 Å resolution cryoEM map describing a complex consisting of full-length mortalin bound to a protein substrate and a full-length GrpEL1 homodimer **(Figure 1D, Supplementary Figure 4)**. To our knowledge, this is the first time human mortalin and human GrpEL1 have been observed in their full-length states and in complex.

**Figure 2:**
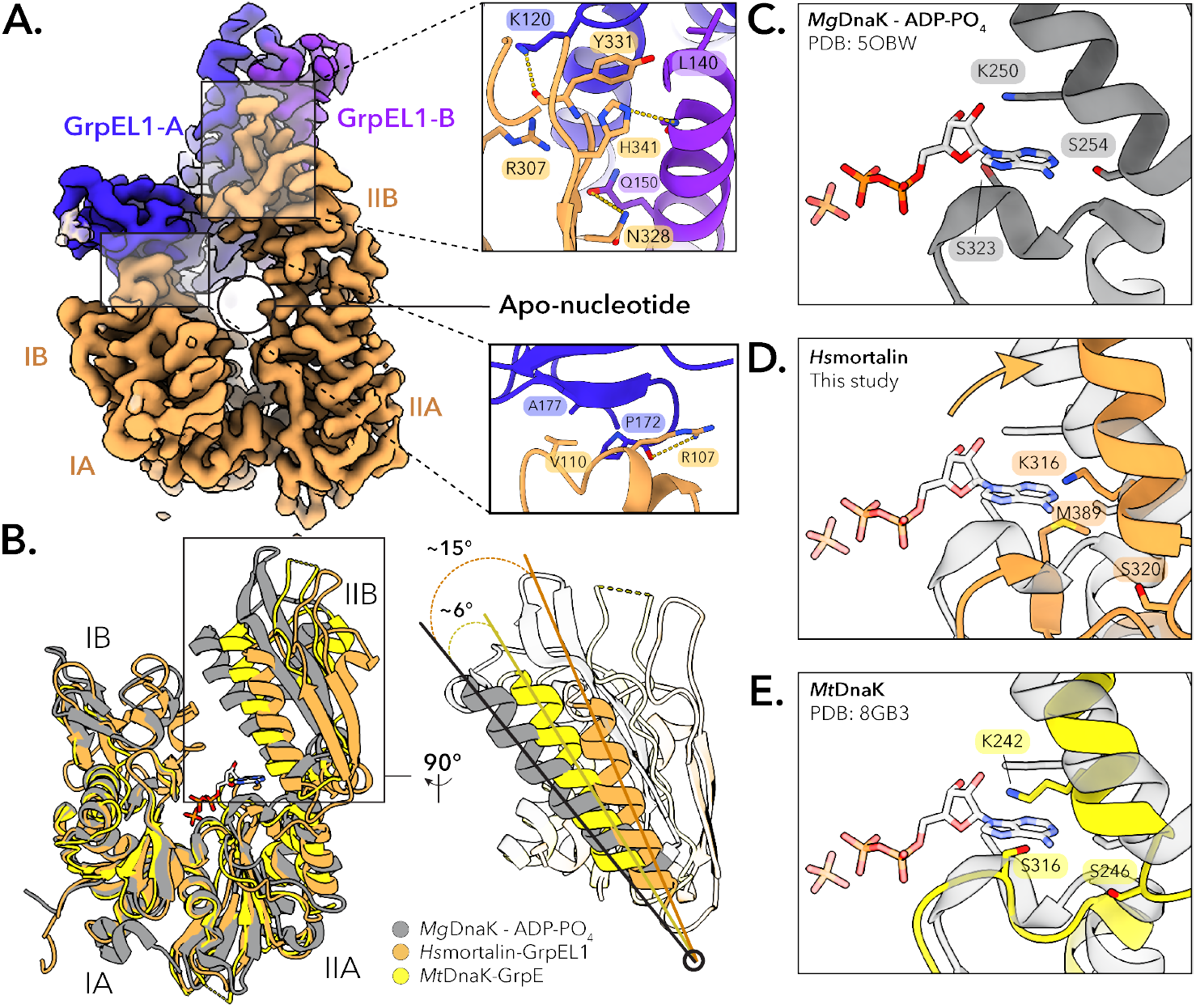
The *Hs*mortalin NBD is fully expanded upon interaction with GrpEL1. **A**. Interface mapping between the mortalin IIB-NBD lobe and the GrpEL1-B short *α*-helix, and the mortalin IB-NBD lobe and the GrpEL1-A *β*-wing domain. Lack of EM density within the NBD suggests an apo-nucleotide NBD. **B**. Structural comparison between the NBDs of *Hs*mortalin-GrpEL1, *Mt*DnaK-GrpE (PDB: 8GB3), and ADP-PO_4_ -bound *Mg*DnaK (PDB: 5OBW). Compared to ADP-PO_4_ -bound *Mg*DnaK, the IIB-NBD lobe of *Hs*mortalin expands ∼15° upon interaction with GrpEL1. **C-E**. Comparison of the ATP binding residues in the IIB-NBD lobe. ATP-stabilizing interactions in the IIB-NBD lobe of *Hs*mortalin-GrpEL1 are removed by movement of the IIB-NBD lobe. ADP-PO_4_ is modeled from *Mg*DnaK-ADP-PO_4_.

**Figure 3:**
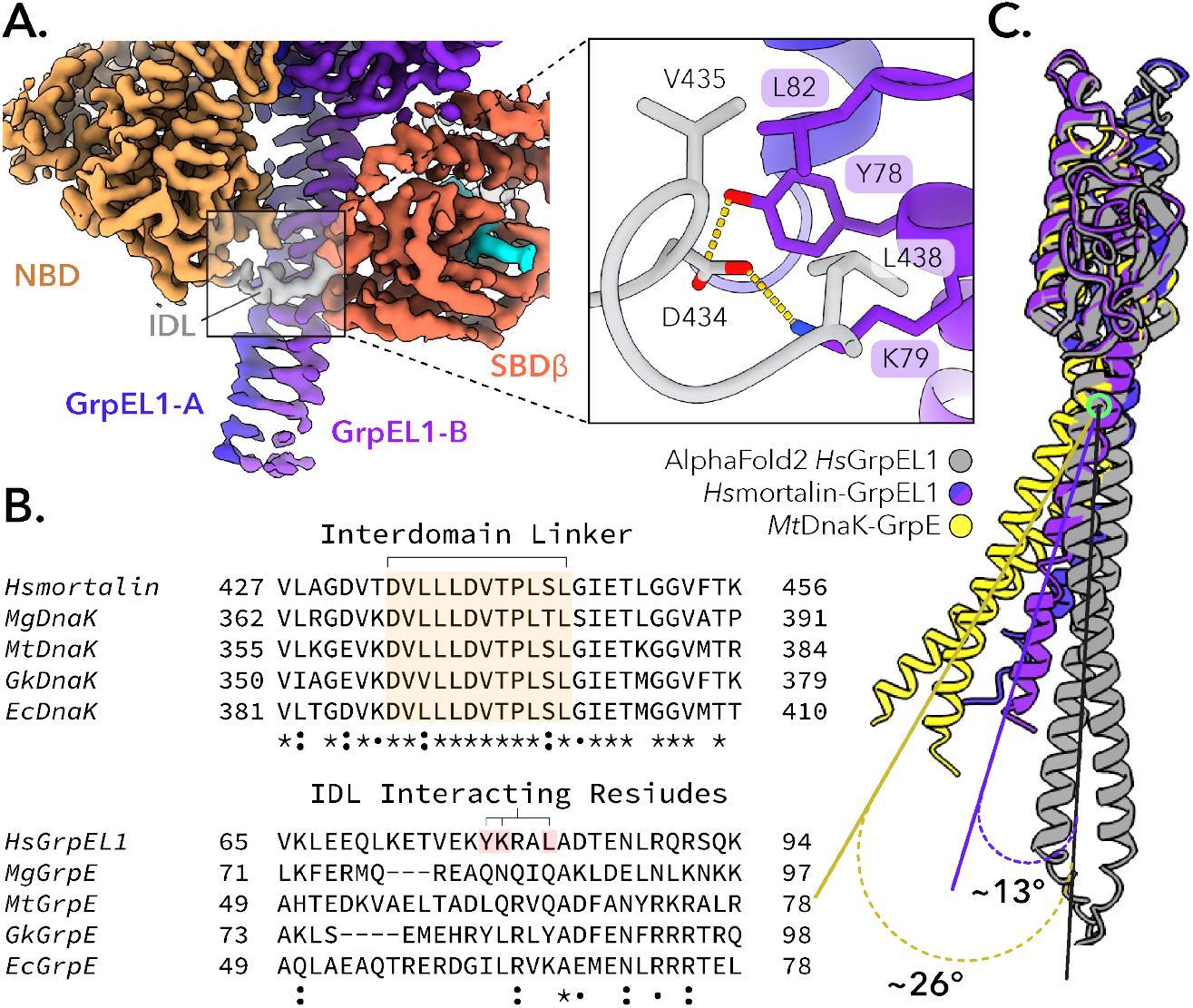
The mortalin interdomain linker is stabilized by interaction with the GrpEL1 long α -helix. **A**. Interaction mapping between the *Hs*mortalin linker and the GrpEL1 long α -helix. **B**. Multiple sequence alignment between Hsp70 and GrpE-like species. The interdomain linker of Hsp70 is highly conserved whereas the interacting GrpE-like region is variable. *Hs*: homo sapiens, *Mg*: mycoplasma genitalium, *Mt*: mycobacterium tuberculosis, *Gk* : geobacillus kaustophilus, *Ec*: eschericia coli. **C**. Superposition of GrpE-like species from our *Hs*mortalin-GrpEL1 structure, *Mt*DnaK-GrpE (PDB: 8GB3), and the AlphaFold2^47^ prediction of *Hs*GrpEL1.

**Figure 4:**
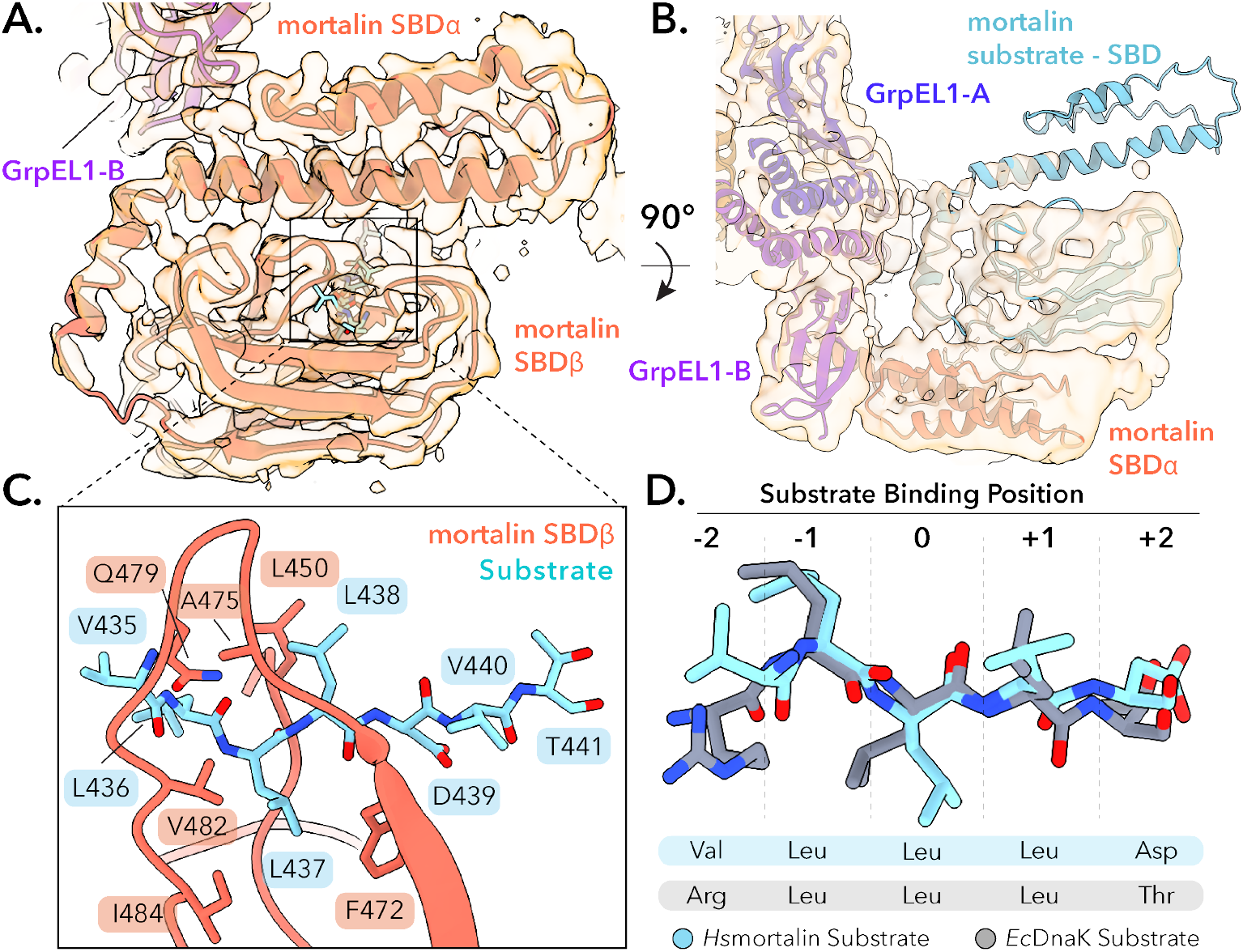
*Hs*mortalin in complex with GrpEL1 is substrate-bound to a mortalin truncation product. **A**. Model of the *Hs*mortalin substrate binding domain (SBD) bound to a substrate fit into the DeepEMhancer-sharpened cryoEM map. **B**. Rigid-body docking of the mortalin SBD into the posterior substrate EM density (low-pass filtered to 5 Å). **C**. Residue mapping of the substrate within the mortalin substrate binding site. **D**. Comparison of the modeled *Hs*mortalin substrate with the crystal structure of *Ec*DnaK bound to the NRLLLTG peptide (PDB: 4EZW).

This cryoEM structure depicts a 1:2 stoichiometry of mortalin:GrpEL1 with mortalin in an elongated state stabilized by numerous interactions across both protomers of the GrpEL1 dimer **(Figure 1D)**. Firstly, we observe an expanded NBD devoid of nucleotide that is mediated by extensive contacts with one protomer of GrpEL1, termed GrpEL1-A. Secondly, we detail significant bending of the GrpEL1 *α*-helical stalk domain that enables interactions between the GrpEL1 stalk and mortalin NBD and SBD. Finally, this structure provides novel insights into the interactions between the SBD*α* helical lid of mortalin and the second GrpEL1 protomer, GrpEL1-B. Critically, lack of EM density within the nucleotide binding pocket but strong density within the substrate binding pocket suggests that our structure represents a previously unobserved intermediate state and suggests a step-wise mechanism for the nucleotide and substrate release action of GrpEL1. Furthermore, since several bacterial DnaK-GrpE structures have been published to date,^22,40,42^ this structure enables novel comparisons between the bacterial and human systems, detailed below.

### GrpEL1 facilitates full expansion of the mortalin NBD for nucleotide release

In our mortalin–GrpEL1 structure, the NBD of mortalin interacts with GrpEL1-A at two distinct interfaces. At the first interface, subdomain IB of the NBD contacts the *β*-wing domain of GrpEL1-A via a salt bridge between R107 in mortalin and the carbonyl group of P172 in GrpEL1-A **(Figure 2A)**. Additional van der Waals interactions between V110 in mortalin and A177 in GrpEL1-A provide additional stabilization to this interface, resulting in 108 Å^2^ of buried surface area. The bulk of the NBD-GrpEL1 interactions, however, are located between the short *α*-helices of GrpEL1-A and GrpEL1-B. At the second interaction interface several electrostatic and van der Waals interactions stabilize the IIB lobe of mortalin’s NBD against GrpEL1-A yielding 219Å^2^ of buried surface area **(Figure 2A)**. Notably, these interfaces are present in bacterial DnaK-GrpE structures and are thought to contribute to the opening of the NBD for nucleotide release.^22,40,42^

Several structures of the bacterial DnaK NBD complexed with GrpE have suggested a nucleotide release mechanism whereby the IIB-NBD lobe rotates outward through interactions with the *β*-wing domain of GrpE, facilitating ADP release.^22,40,42^ We observe a similar expansion of the mortalin NBD in our structure, as well as a lack of EM density within the nucleotide binding pocket **(Figure 2A)**. To probe the potential differences between the human and bacterial nucleotide release mechanisms, we performed structural comparisons between various DnaK-GrpE structures and our mortalin–GrpEL1 structure. Aligning these structures to the NBD IA and IB lobes allowed for visualization of the motions associated with GrpEL1-mediated rotations of the IIB lobe. Using the inward-facing *α*-helix within the IIB lobe as a point of reference, our structure exhibits ∼15° of rotation compared to the ATP-bound *Mg* DnaK structure (PDB: 5OBW)^43^ and ∼6° of increased rotation compared to the *Mt* DnaK-GrpE structure (PDB: 8BG3)^40^ **(Figure 2B)**. Compared to existing mortalin NBD structures (PDB: 4KBO, 6NHK)^12,31^ we observe a similar widening of the IIB lobe **(Supplementary Figure 5)**. Together, these analyses indicate conserved mechanisms of GrpE(L1)-mediated ADP release from the (mt)Hsp70 NBD across kingdoms.

**Figure 5:**
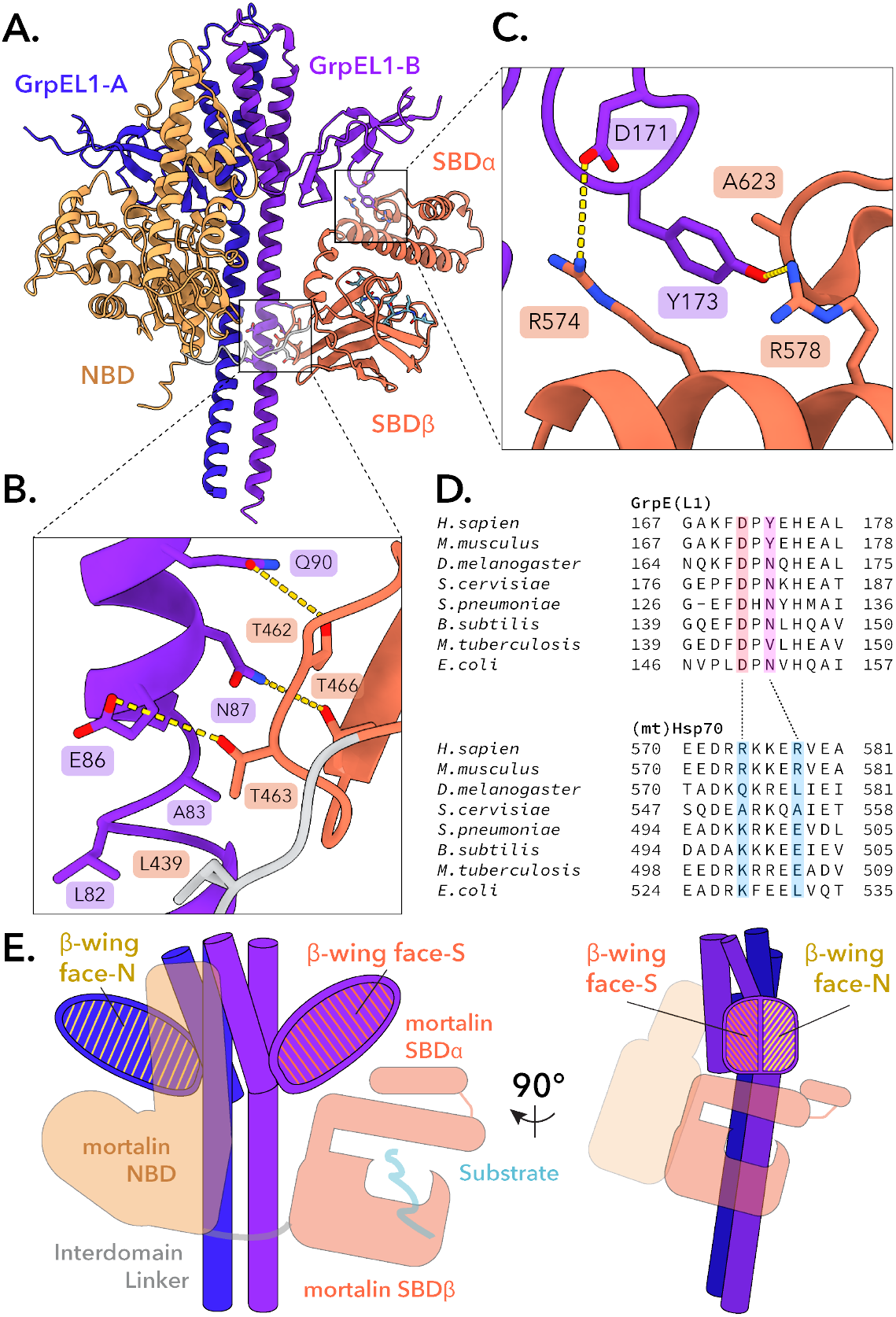
GrpEL1 forms unique interactions with the *Hs*mortalin NBD and SBD. **A**. Structural mapping of the *Hs*mortalin SBD with the GrpEL1-B long α -helix and *β*-wing domain. **B**. Residue mapping of the interactions between the GrpEL1-B long α -helix and *Hs*mortalin SBD*β* cleft. **C**. Residue mapping of the interactions between the GrpEL1-B *β*-wing domain and SBD*α* helical domain. **D**. Multiple sequence alignment of the GrpEL1-B *β*-wing-SBD*α* interacting regions across GrpE-like and Hsp70 species. **E**. Designation of the GrpEL1 *β*-wing face that interacts with the *Hs*mortalin NBD (*β*-wing face-N) and the face that interacts with the SBD (*β*-wing face-S).

Several conserved residues in the Hsp70 nucleotide binding pocket have been identified to stabilize bound nucleotide.^6,44–46^ In our structure, nucleotide-interacting residues residing in the IA-NBD lobe are in similar positions relative to the ATP-bound state. However, residues K316 and S320 (K250 and S254 in *Mg* DnaK)^43^ are displaced, abolishing stabilizing interactions of the ribose and adenosine rings of a bound nucleotide **(Figure 2C-E)**. This suggests that displacement of key residues in the IIB lobe destabilize the bound nucleotide and contribute to ADP release.

### The mortalin interdomain linker facilitates bending of the GrpEL1 stalk

In our mortalin–GrpEL1 structure we observe EM density for the interdomain linker (IDL) region that spans the nucleotide and substrate binding domains. Here, the mortalin IDL is proximal to the GrpEL1 *α*-helical dimer with residues V435 and L438 within the hydrophobic IDL interacting with L82 in GrpEL1-B **(Figure 3A)**. Electrostatic and hydrogen bonding interactions between D434 in the IDL and K79 and Y78 in GrpEL1-B further stabilize the mortalin IDL **(Figure 3A)**, contributing 146 Å^2^ of surface area. Previous structures of bacterial DnaK bound to GrpE have described a similar IDL-GrpE interaction.^31,40,42^ To further evaluate the significance of this interaction, we compared the residues of this interacting region between human and bacterial Hsp70-GrpE systems. Interestingly, while the IDL region is highly conserved across human and bacterial sequences, the corresponding interacting region within GrpE-like species varies significantly **(Figure 3B)**.

Interaction between the mortalin IDL and GrpEL1 appears to be facilitated by significant bending of the GrpEL1 α -helical domain. Previous structures of bacterial GrpE and DnaK-GrpE that contain the IDL of DnaK describe a similar bending of the GrpEL1 stalk but to varying magnitudes.^31,40,42^ To quantify the degree of GrpEL1 bending we compared our structure and other DnaK-GrpE structures to the AlphaFold2^47^ prediction of a human GrpEL1 dimer, which describes a linear α -helical domain. Compared to the predicted GrpEL1 structure, our mortalin-GrpEL1 structure exhibits a ∼13° bending of the GrpEL1 *α*-helical domain. In contrast, the structure of MtDnaK-GrpE^40^ exhibits a ∼26° bending of the α -helical domain **(Figure 3C)**.

### Substrate interactions within the mortalin–GrpEL1 complex

Our EM density reveals the entirety of the mortalin substrate binding domain, with the SBD α -helical subdomain resting on top of the *β*-sandwich subdomain **(Figure 4A)**. We observed strong density for a peptide bound to the SBD substrate binding pocket, indicating that our purified complex is substrate-bound. Behind this clear substrate density, we observed weaker density that matches the mortalin SBD *β*-sandwich subdomain **(Figure 4B)**. Hsp70s have been observed to bind themselves as substrates as in the *Gk* DnaK-GrpE (PDB: 4ANI)^42^ crystal complex. Thus, we hypothesized that the mortalin protomer in complex with GrpEL1 was bound to the interdomain linker region of another mortalin protomer. The high resolution of our structure enabled modeling of the bound peptide as VLLLDVT, corresponding to residues 435-441 of the mortalin IDL. We did not observe density for the NBD of the second mortalin protomer. Given the stability of the NBD, it is unlikely that the absence of NBD density can simply be attributed to signal averaging. Rather, we propose that the observed bound substrate is a mortalin truncation product containing only the IDL and SBD. We performed size exclusion chromatography coupled with multi-angle light scattering (SEC-MALS) on our mortalin-GrpEL1 complex which indicated the presence of a 152.5 kDa species **(Supplementary Figure 2)**. This molecular weight agrees with a complex consisting of one full-length mortalin (70.2 kDa), two GrpEL1 protomers (25.5 kDa each) and an IDL-SBD mortalin truncation product (28.5 kDa). Additionally, modelling of the mortalin NBD connected to an IDL bound to the substrate binding pocket appears to be capable of accommodating a full-length mortalin **(Supplementary Figure 6)**. Taken together, these observations suggest that our mortalin-GrpEL1 complex is bound to a second partial mortalin protomer as a substrate.

**Figure 6:**
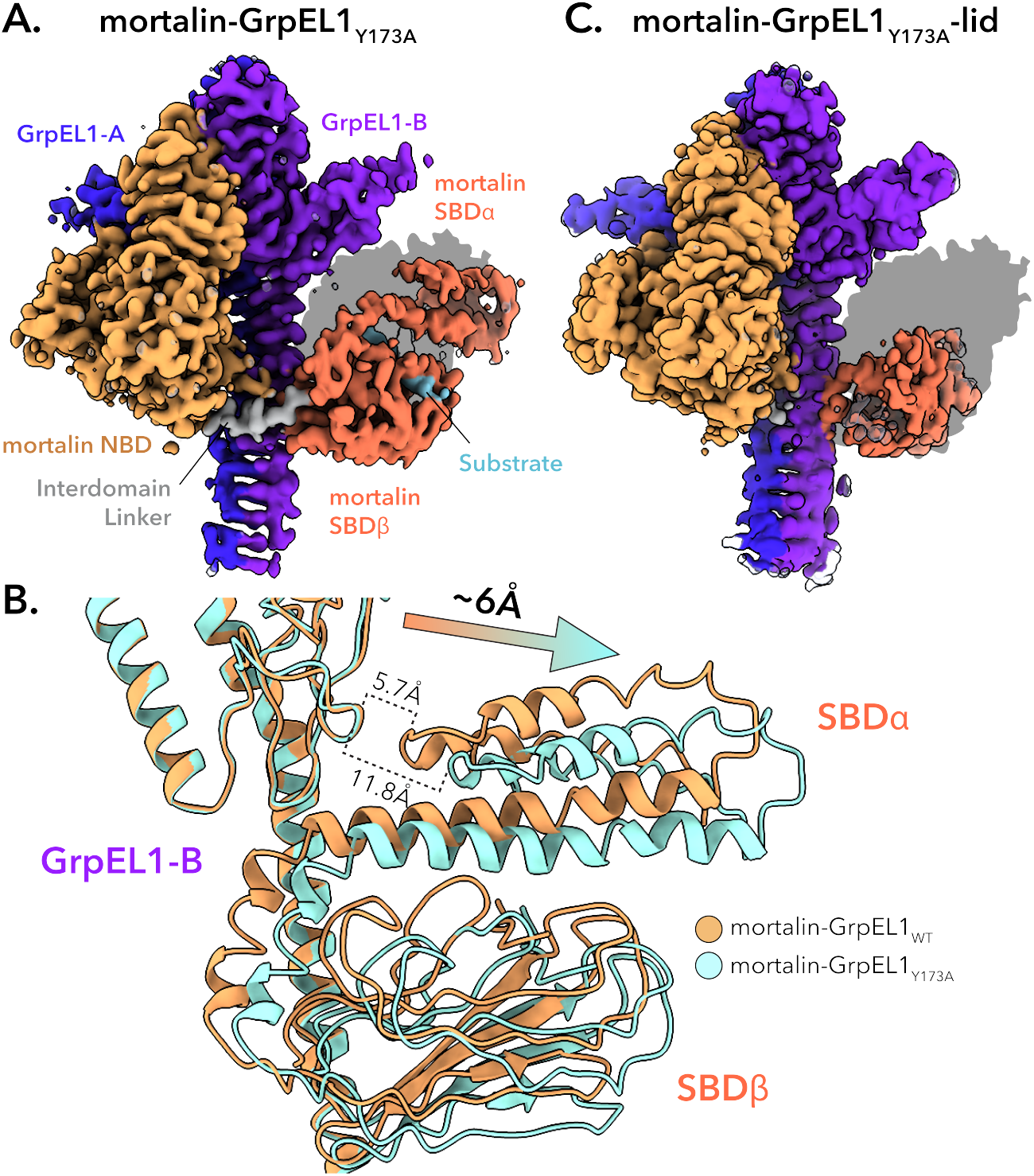
Mutation of Y173A in GrpEL1 results in a shift of the *Hs*mortalin SBD. **A**. DeepEMhancer-sharpened map of mortalin-GrpEL1_*Y* 173 *A*_. Shadow represents the silhouette of the mortalin-GrpEL1_*WT*_ SBD position. **B**. Superposition of mortalin-GrpEL1_*WT*_ and mortalin-GrpEL1_*Y* 173 *A*_. In the mortalin-GrpEL1_*Y* 173 *A*_ structure, the SBD is translated ∼6 Å away from GrpEL1-B compared to WT. **C**. DeepEMhancer-sharpened map of mortalin-GrpEL1_*Y* 173 *A*_ -lid. Shadow represents the silhouette of the mortalin-GrpEL1_*WT*_ SBD position.

Hsp70 substrate specificity and binding modes have been studied extensively using various biochemical and structural techniques.^5,6,48,49^ Current Hsp70 binding models are based upon DnaK-substrate interactions and describe a hydrophobic binding pocket that can accommodate five to seven hydrophobic residues.^5,6,48,49^ The central residue, termed the 0th position, favors the binding of leucine residues whereas the flanking residues are typically occupied by hydrophobic residues.^5,49^ In our structure we identify a mortalin substrate binding mode consistent with the canonical DnaK substrate binding model. Here, a leucine is oriented in the 0th position and makes hydrophobic contacts with I484 and F472.

Adjacent to the 0th leucine are additional flanking leucine residues which make additional hydrophobic contacts with L450, A475, and V482 **(Figure 4C)**. Comparison between the mortalin substrate and an *Ec*DnaK substrate (PDB: 4EZW)^50^ displays a highly conserved binding mode with several central leucines forming the core of the peptide substrate **(Figure 4D)**. Overall, our structure suggests that mortalin substrate binding occurs, as expected, in a near identical mode compared to DnaK substrate binding.

### Interactions between GrpEL1-B and the SBD domain of mortalin

In our structure, the GrpEL1-B protomer interacts with both of mortalin’s SBD subdomains **(Figure 5A)**. First, downstream of mortalin’s inter-domain linker region several electrostatic and hydrogen bonding interactions between the SBD*β* cleft and the N-terminal stalk of GrpEL1-B result in 180 Å ^2^ of buried surface area **(Figure 5B)**. Second, we observe a new interaction interface between the SBD α helical domain and the *β*-wing domain of GrpEL1-B. Here, Y173 from GrpEL1-B inserts itself into the mortalin SBD*α*, forming electrostatic and cation-pi stacking interactions with R578 and R574, respectively. Additionally, D171 in GrpEL1-B forms an electrostatic interaction with R574 to form an interface with a total buried surface area of 159 Å^2^ **(Figure 5C)**. To interrogate the prevalence of this interface, we performed a co-variational analysis between Hsp70 and GrpE homologs of residues at this interface.

First, we compared the electrostatic interaction between D171 in *Hs*GrpEL1 and R574 *Hs*mortalin across species. In all sequences compared, ranging from bacteria to higher eukaryotes, D171 (*Hs*GrpEL1) was highly conserved **(Figure 5D, Supplementary Figure 7)**. Interestingly, R574 (*Hs*mortalin) corresponded to mostly arginines and lysines in other species which would allow preservation of the salt bridge spanning the GrpEL1-B *β*-wing and SBD*α* lid. In our structure, Y173 (*Hs*GrpEL1) appears buried within the SBD α lid and participates in electrostatic, cation-pi, and hydrophobic interactions with mortalin. While this tyrosine is highly conserved in vertebrates, this residue is substituted for mostly Asn in bacteria and lower eukaryotes. Interestingly, residues corresponding to R578 (*Hs*mortalin) are substituted for negatively charged (Asp/Glu) and hydrophobic (Leu) residues.

**Figure 7:**
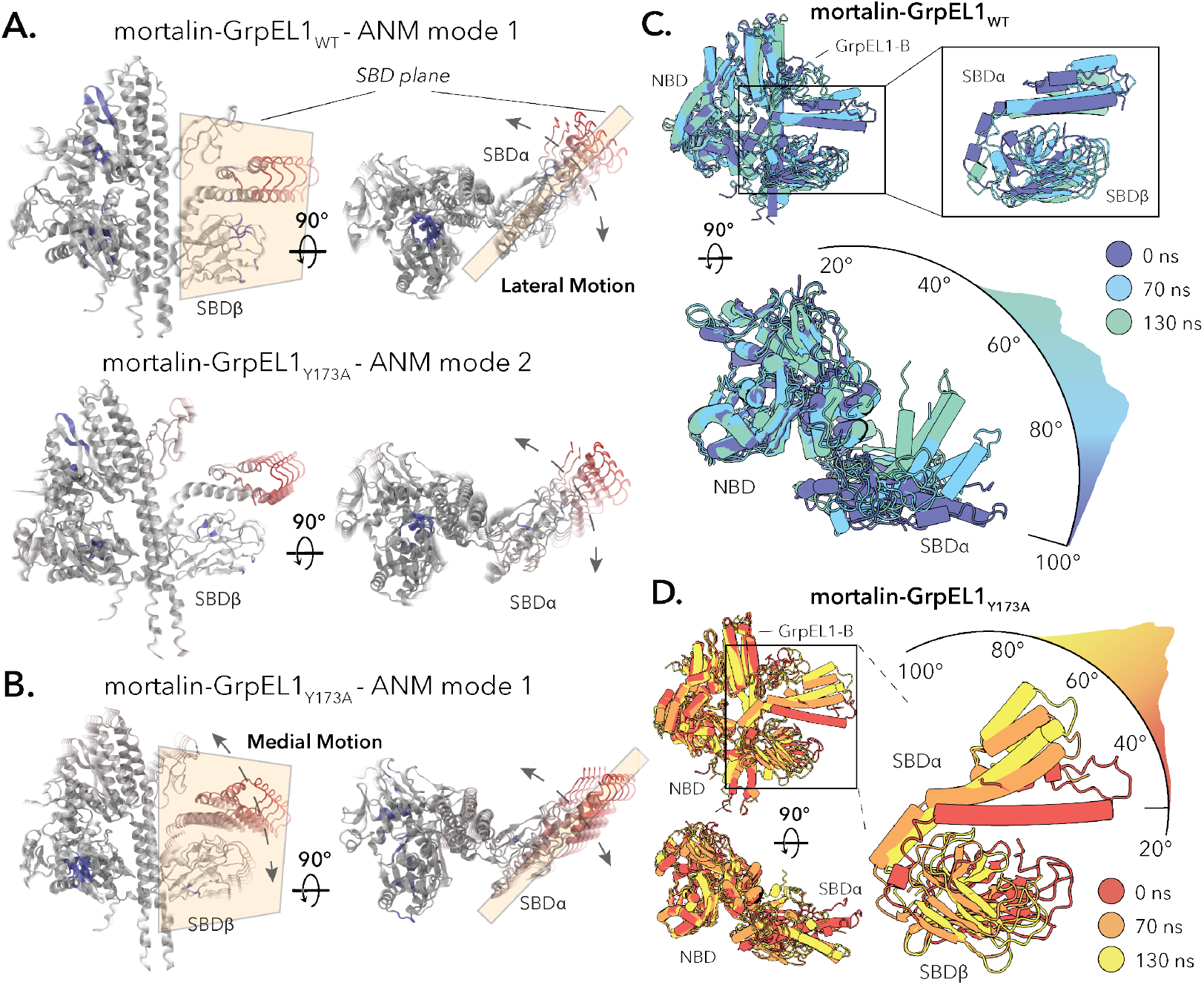
Flexibility analysis of mortalin-GrpEL1_*WT*_ and mortalin-GrpEL1_*Y* 173 *A*_. **A**. Anisotropic network modeling (ANM) modes of the SBD*α* lateral motion observed in both mortalin-GrpEL1_*WT*_ and mortalin-GrpEL1_*Y* 173 *A*_. SBD plane is colored in beige. **B**. ANM mode of the lateral and medial SBD α motion found exclusively in the mortalin-GrpEL1_*Y* 173 *A*_ ANM analysis. SBD plane is colored in beige. **C**. All-atom MD simulation of mortalin-GrpEL1_*WT*_. The position of the SBD α lid lateral motion is visualized across a representative 150ns simulation. Timepoints at 0, 70, and 130ns are represented as the start, middle, and end of the simulation. **D**. All-atom MD simulation of mortalin-GrpEL1_*Y* 173 *A*_. The medial motion of the SBD*α* lid is visualized across the 150ns simulation. Timepoints at 0, 70, and 130ns are represented as the start, middle, and end of the simulation.

Negatively charged residues would be capable of interacting with the substituted Asn and maintain this GrpE *β*-wing-SBD α interface **(Supplementary Figure 8)**.

**Figure 8:**
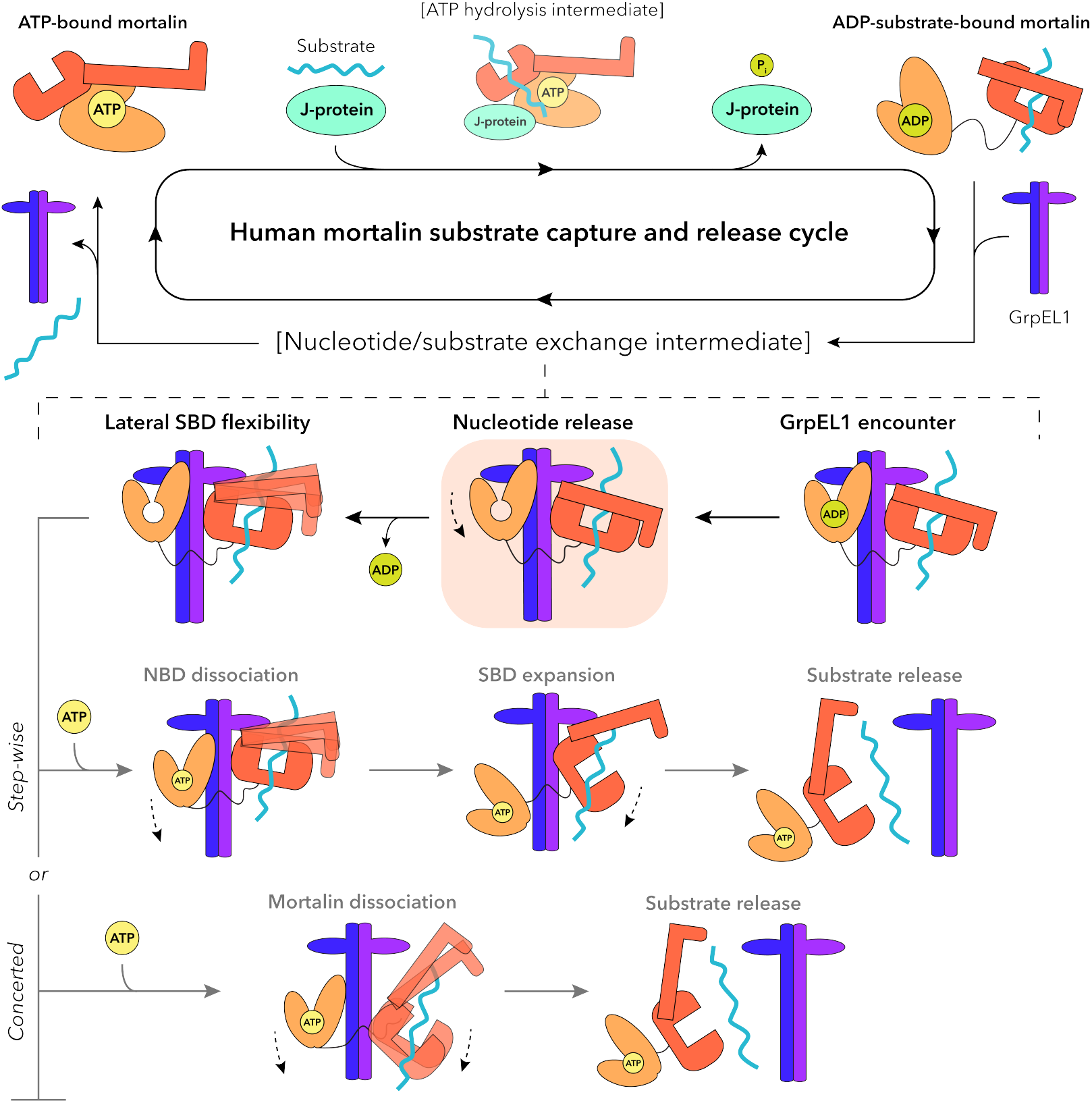
GrpEL1-mediated nucleotide and substrate release mechanism of human mortalin. Upon interaction with a substrate and J-protein, mortalin hydrolyzes ATP and stably interacts with substrate. GrpEL1 interacts asymmetrically with mortalin and facilitates ADP release via interactions at the NBD. Flexibility within the SBD loosens the SBD α subdomain. Following ATP binding in a step-wise mechanism, the NBD dissociates first which enables opening of the SBD and subsequent substrate release. Alternatively, ATP binding induces concerted dissociation of mortalin whereby decoupling of SBD α and GrpEL1-B enables substrate release. The intermediate representing the mortalin-GrpEL1_*WT*_ structure is highlighted in orange.

The interactions we observe between the GrpEL1-B *β*-wing and SBD*α* are exclusive to this interface since Y173 in GrpEL1-A does not appear to contact the NBD **(Supplementary Figure 9)**. Considering the unique interaction interfaces we observe between NBD-GrpEL1-A and SBD-GrpEL1-B, we designate unique identifiers to the GrpEL1 faces that interact with the NBD, termed *β*-wing face-N, or interact with the SBD, termed *β*-wing face-S **(Figure 5E)**. These unique interaction faces rationalize the necessity of a dimeric GrpEL1 and support the specific, asymmetric mortalin conformation with GrpEL1. Intrigued by the GrpEL1-B-SBD*α* interface, we hypothesized that interactions here contribute to GrpEL1-mediated substrate release. Thus, we mutated Y173 in GrpEL1 to an alanine (GrpEL1_*Y* 173 *A*_) and performed cryoEM analysis on a mortalin-GrpEL1_*Y* 173 *A*_ complex.

### CryoEM structure determination of mortalin-GrpEL1_*Y* 173 *A*_

Complex formation with mortalin and GrpEL1_*Y* 173 *A*_ yielded a similar profile on size exclusion chromatography compared to mortalin-GrpEL1_*WT*_ **(Supplementary Figure 10)**. As previously described, the high molecular weight peak was concentrated and vitrified onto cryoEM grids. Our cryoEM analysis of mortalin-GrpEL1_*Y* 173 *A*_ yielded two distinct populations differentiated by the presence or lack of density for the SBD*α* lid. We term these structures mortalin-GrpEL1_*Y* 173 *A*_ and mortalin-GrpEL1_*Y* 173 *A*_ -lid **(Supplementary Figures 11-12)**. Overall, these structures closely resemble our mortalin-GrpEL1_*WT*_ structure and are similarly nucleotide-free. However, we observed notable differences in the IDL and SBD regions of our mortalin-GrpEL1_*Y* 173 *A*_ structures. Firstly, our mortalin-GrpEL1_*Y* 173 *A*_ structure harbors EM density for the entirety of the SBD, and similar to our mortalin-GrpEL1 structure, appears substrate bound to another mortalin proteolysis product **(Figure 6A)**. Critically, we observe a large deviation of the SBD in our mortalin-GrpEL1_*Y* 173 *A*_ structure compared to mortalin-GrpEL1_*WT*_, with the SBD of mortalin-GrpEL1_*Y* 173 *A*_ translated ∼6 Å away from the GrpEL1-B *β*-wing **(Figure 6B)**. In our mortalin-GrpEL1_*Y* 173 *A*_ -lid structure, we discern diminished EM density for the IDL region and density only for the SBD*β* subdomain **(Figure 6C)**. Importantly, we are unable to identify EM density of bound substrate that is observed in our other mortalin-GrpEL1 structures. Considering density for the SBD α subdomain is also absent in the mortalin-GrpEL1_*Y* 173 *A*_ -lid structure, we hypothesize that this com-plex harbors a disordered SBD α subdomain and may represent a post-substrate release state. A recently published structure of *Mt* DnaK-GrpE (PDB: 8GB3)^40^ also harbors a disordered SBD α subdomain and is suggested to represent a post-substrate release conformation.

These findings are intriguing for several reasons. Firstly, positioning of the SBD in our mortalin-GrpEL1_*Y* 173 *A*_ structure suggests that although Y173 in GrpEL1 is not necessary for mortalin-GrpEL1 complex formation, it is key in mediating interaction between the *β*-wing domain of GrpEL1-B and SBD α lid of mortalin. Secondly, presence of the full SBD in mortalin-GrpEL1_*Y* 173 *A*_ suggests that interactions between the GrpEL1-B stalk and SBD*β* subdomain of mortalin are sufficient in stabilizing the entirety of the SBD. Finally, lack of SBD α density in our mortalin-GrpEL1_*Y* 173 *A*_ -lid structure suggests flexibility of the SBD α lid, especially in the absence of substrate.

### Mortalin-GrpEL1 is dynamic and exhibits large movements in the SBD α lid

To investigate the potential flexibility of our structures, we performed anisotropic network modeling (ANM) of mortalin-GrpEL1_*WT*_ and mortalin-GrpEL1_*Y* 173 *A*_ **(Supplementary Videos 1 & 2)**. Mode 1 of the mortalin-GrpEL1_*WT*_ analysis shows a large motion in the SBD α sub-domain that is perpendicular to the plane formed by SBD α and SBD*β* that we term a lateral motion. **(Figure 7A**,**top)**. Mode 2 of our mortalin-GrpEL1_*Y* 173 *A*_ analysis also shows a similar lateral motion **(Figure 7A, bottom)**. Interestingly, mode 1 in mortalin-GrpEL1_*Y* 173*A*_ describes a prominent diagonal motion in the SBD*α* lid that appears to be a combination of the observed lateral motion and another medial motion whereby the SBD α moves parallel to the SBD α /*β* plane **(Figure 7B)**.

To validate the motions observed in our ANM analyses, we conducted 450 nanoseconds of all-atom molecular dynamics (MD) simulations on the mortalin-GrpEL1_*WT*_ and mortalin-GrpEL1_*Y* 173 *A*_ structures (three replicates of 150 nanoseconds each), sufficient to capture domain-level motions in the complex. In our mortalin-GrpEL1_*WT*_ simulations, we observed the large lateral motion of the SBDα lid observed in mode 1 of the ANM analysis **(Figure 7C)**. We found that the SBD*α* rotated up to 50° from its starting position and spent significant time at 20-40° throughout the course of the simulation **(Figure 7C, Supplementary Figure 13)**. Comparisons between the SBD in mortalin-GrpEL1_*WT*_ and existing Hsp70-SBD structures reveal positional variations of the SBD α lid, providing experimental evidence of lateral flexibility in the SBD α subdomain **(Supplementary Figure 14)**. In simulations with mortalin-GrpEL1_*Y* 173 *A*_, we observed a prominent medial motion parallel to the SBD plane where the SBD*α* lid extends up to ∼35° away from SBD*β*, exposing the substrate binding pocket **(Figure 7D)**. Throughout the simulation, the SBD α spent most of the time 20-35° away from its initial position **(Figure 7D)**. Notably, while two of the three mortalin-GrpEL1_*Y* 173 *A*_ simulations described this medial motion, one simulation described a combination of lateral and medial motions remin-iscent of mode 1 from the ANM analysis **(Supplementary Figure 15)**. Taken together, our ANM analyses and all-atom MD simulations indicate that the SBD*α* lid is highly dynamic. Importantly, we observe distinct motions associated with either mortalin-GrpEL1_*WT*_ or mortalin-GrpEL1_*Y* 173 *A*_, where the WT GrpEL1 complex exhibits a dominant lateral motion of the SBD αlid, and the Y173A GrpEL1 complex exhibits a combination of lateral and medial motions. These analyses underscore the dynamic nature of the SBD α subdomain and how uncoupling of the SBD α -GrpEL1-B interface endows greater flexibility to the SBD. Finally, these motions point towards a potential substrate release mechanism where GrpEL1-mediated flexibility in the SBD may prime mortalin for substrate release.

## Discussion

We used cryoEM to investigate how human GrpEL1 facilitates nucleotide exchange and substrate release of human mortalin. Our structure describes a 1:2 stoichiometry between mortalin and GrpEL1 that is analogous to previously observed bacterial DnaK-GrpE structures.^22,40,42^ Importantly, we do not observe nucleotide in the NBD but strong density within the SBD, suggesting GrpEL1 facilitates nucleotide exchange prior to substrate release. We further identify a novel interface between mortalin-GrpEL1 and a key interaction stabilizing a distinct mortalin SBD conformation. Together, our analyses corroborate over two decades of work on Hsp70-GrpE complexes and enable us to speculate on the mechanisms underlying GrpEL1-mediated substrate release.

In our cryoEM structure of full-length *Hs*mortalin-GrpEL1 we observe a significant separation of the NBD lobes mediated by GrpEL1-A’s *β*-wing domain. Compared to bacterial DnaK-GrpE structures,^22,40,42^ our complex shows a larger rotation in the IIB-NBD lobe. Considering the high sequence similarity between mortalin-GrpEL1 and DnaK-GrpE, it is unlikely that this difference is solely attributed to human vs. bacterial systems. Instead, we propose that docking of the SBD to the GrpEL1-B *β*-wing domain enables full expansion of the NBD, minimizing the chances of pre mature substrate release. Additionally, we note a moderate (∼13°) bending of the GrpEL1 long α-helical stalk that is likely constrained by myriad interactions across mortalin-GrpEL1. Finally, our identification of *β*-wing face-N and *β*-wing face-S underscores the importance of an asymmetric GrpEL1 homodimer and rationalizes the 1:2 stoichiometry of Hsp70:GrpE-like structures.

The interaction between GrpEL1-B and the SBD in our mortalin-GrpEL1 complex provides new insights that were lacking in previous DnaK-GrpE homolog structures. We observe clear EM density spanning the GrpEL1-B *β*-wing domain and SBD α lid subdomain, indicative of a stable interface. Mutational and structural analyses of the mortalin-

GrpEL1_*Y* 173 *A*_ complex highlight the crucial role of Y173 in maintaining connectivity at this interface. However, the presence of an intact, substrate-bound SBD in our mortalin-GrpEL1_*Y* 173 *A*_ structure suggests that interactions between SBD*β* and GrpEL1-B stalk are sufficient to maintain the closed SBD conformation proximal to GrpEL1.

In our study we used a mutant construct of mortalin harboring the R126W point mutation. This mutation has been implicated in EVEN-PLUS syndrome and exhibits decreased ATPase activity and stability.^12^ In mortalin-GrpEL1_*WT*_ we do not observe R126W making additional contacts within the NBD **(Supplementary Figure 16A)**. In the *Mt* DnaK-GrpE structure (PDB: 8GB3),^40^ the corresponding S75 in *Mycobacterium tuberculosis* does not appear to contribute to complex stabilization, supporting the idea that R126 does not contribute to mortalin-GrpEL1 complex formation **(Supplementary Figure 16B)**. Interestingly, upon examination of the AlphaFold2^47^ predicted structure of human mortalin, R126 appears to form contacts with T271 in NBD lobe II **(Supplementary Figure 16C)**. Thus, it is possible that R126 is important for stabilization of a closed NBD amenable for ATP binding.

Our analyses provide novel insights into the architecture and dynamics of *Hs*mortalin-GrpEL1 and enable us to propose a new mechanism of GrpEL1-mediated regulation of mortalin. Here, ADP-bound mortalin with substrate encounters GrpEL1 and adopts an asymmetric conformation with the NBD interacting with *β*-wing face-N of GrpEL1-A, and SBD α interacting with *β*-wing face-S of GrpEL1-B. GrpEL1-A interactions with the IIB-NBD lobe result in expansion of the NBD and release of ADP. Lateral flexibility enables loosening of the SBD α lid, potentially priming the complex for subsequent substrate release. Following ATP binding to the NBD we propose two potential pathways towards substrate release and complex dissociation. In the step-wise pathway, the NBD dissociates from GrpEL1 first and pulls away from SBD*β*. The SBD α lid remains in contact with GrpEL1-B as SBD*β* opens and enables substrate release, followed by full complex dissociation. In an alternative pathway, dissociation of the NBD and SBD are concerted, and substrate is released by flexibility of the SBD upon decoupling of the SBD α lid-GrpEL1-B interface **(Figure 8)**. Further studies of the allosteric changes induced by ATP addition will be required to identify a comprehensive substrate release mechanism.

Several biochemical studies have demonstrated the importance of the disordered N-terminal region of GrpE-like proteins in facilitating substrate release. However, it should be noted that these studies have been limited to bacterial DnaK-GrpE systems and given the substantial sequence divergence of the GrpEL1 N-terminus, partly due to the presence of a mitochondrial targeting sequence, we cannot assume conservation of this mechanism in eukaryotes. Nevertheless, we can speculate on the functional implications of eukaryotic GrpEL1 isoforms. Vertebrates possess two mitochondrially targeted GrpE-like isoforms, GrpEL1 and GrpEL2, with GrpEL2 non-essential but up-regulated under oxidative stress.^51–53^ Notably, substitution of Y173 with H176 in GrpEL2 may suggest an additional level of regulation in stress-responsive exchange factors **(Supplementary Figure 17)**.

Together, our work reveals a near-complete molecular mechanism for GrpEL1 regulation of human mortalin. Additional investigations into how ATP mechanistically induces substrate release and complex dissociation will be key to obtain a comprehensive understanding of this process. Importantly, our cryoEM structures represent the first visualization of full-length, human mortalin and GrpEL1 and shed light on fundamental Hsp70 mechanisms ubiquitous to biology. These insights will be key in future studies to understand cellular responses in basal and stressed environments, and in the development of pharmaceuticals targeting Hsp70-related diseases.

## Methods

### Plasmid/Construct Design

Plasmids encoding full-length genes for *Hs*Hep1 and *Hs*mortalin (C-terminal hexahistidine tag) in a pETDuet-1 vector, human GrpEL1 (C-terminal tandem Strep-Tag II and hexahistidine tags) in a pRSFDuet-1 vector, as well as human Pam16 and Pam18 in a pCDFDuet-1 vector were obtained from Bio-Basic. The mitochondrial targeting sequences for each gene (residues 1-49 for Hep1, 1-46 for mortalin, and 1-27 of GrpEL1) were subsequently removed prior to recombinant expression in *Escherichia coli*. The R126W mutant of mortalin (hereafter referred to as mortalin) and Y173A mutant of GrpEL1 (GrpEL1_*Y* 173*A*_) were generated using around-the-horn PCR. We designed a chimeric form of the peripheral membrane proteins Pam16 and Pam18 to eliminate the transmembrane helices and improve solubility of the J-like/J-protein heterodimeric Pam16/18 complex. Briefly, we fused the functional J-like (residues 56-121) and J-domains (residues 48-116) of *Hs*Pam16 and *Hs*Pam18, respectively, to the N- and C-terminus of *E. coli* maltose binding protein (MBP), respectively, with a 3x(GGS) linker between Pam16 and MBP and a GGS-HHHHHH-GGS linker between MBP and Pam18. We used AlphaFold2^47^ to predict the structure of our Pam16(J-like)-MBP-H6-Pam18(J-domain) (Pam16/Pam18 chimera) chimeric complex and observed an architecture similar to that of the yeast Pam16/Pam18 crystal structure (PDB ID: 2GUZ), suggesting our chimeric complex could be functional **(see Supplementary Figure 1)**. All oligonucleotides were ordered from Integrated DNA Technologies and all plasmid sequences were confirmed by GENEWIZ. Verified plasmid sequences were transformed into the *E*.*coli* LOBSTR^55^ strain for recombinant protein expression.

### Recombinant protein expression and purification

#### Mortalin

A pETDuet-1 vector encoding for *Hs*Hep1 and *Hs*mortalin was transformed into the *E*.*coli* LOBSTR^55^ strain and grown in LB medium containing 50*µ*g/mL ampicillin and 50*µ*M ZnCl_2_ at 37°C until the OD600 reached 1.0. Protein expression was induced by addition of 100*µ*M isopropyl *β*-D-1-thiogalactopyranoside (IPTG) and allowed to express for 4 hrs at 16°C. Cells were centrifuged, collected, and resuspended in buffer A (20mM Tris pH 8, 150mM KCl, 50*µ*M ZnCl_2_, 5mM *β*-mercaptoethanol (BME), 25mM imidazole, 1mM MgCl_2_, 1mM CaCl2, 1*µ*g/mL polyethyleneimine (PEI), 1mM phenylmethylsulfonyl fluoride (PMSF), 5mM benzamidine, 10*µ*M leupeptin, 1*µ*M pepstatin A, and 2*µ*g/mL aprotinin) with a small amount of DNase I (Millapore Sigma). Cells were lysed using a Branson SFX550 sonifier on ice. Lysed cells were centrifuged at 20,000 x g for 30 min at 4°C and the clarified supernatant was loaded onto a 5mL HisTrap HP column (Cytiva) pre-equilibrated with buffer A at 1mL/min using a peristaltic pump. The column was washed with buffer B (20mM Tris pH 8, 150mM KCl, 50*µ*M ZnCl_2_, 5mM BME, and 100mM imidazole) until protein was no longer observed in the eluent, as visualized by mixing with Bradford reagent. The target protein was eluted using 5 column volumes (CVs) of buffer B containing 300mM imidazole. Collected elutions were incubated for 30 min on ice with 0.5mM AMP-PNP and 2mM MgCl_2_ prior to mixing with buffer C (20mM Tris pH 8, 5mM BME) to lower the [KCl] to 75mM. A 5mL HiTrap Q HP anion exchange chromatography column (Cytiva) was pre-equilibrated with buffer C supplemented with 75mM KCl (Q buffer A). Mortalin fractions were loaded onto the HiTrap Q HP column at 1 mL/min and washed with Q buffer A until baseline was reached. A linear gradient was applied using Q buffer A to Q buffer B (20mM Tris pH 8, 1M KCl, 5mM BME) from 0-100% Q buffer B over 20 CVs. Peak fractions containing mortalin were pooled, concentrated and applied to a Superdex 200 Increase 10/300 GL (Cytiva) that had been pre-equilibrated with buffer D (20mM Tris pH 8, 150mM KCl, 0.5mM TCEP). Peak fractions were collected and analyzed using SDS-PAGE. The cleanest fractions were combined and concentrated to 1mg/mL. Concentrated fractions were used immediately, or flash frozen for storage.

#### GrpEL1_WT_

A pRSFDuet-1 vector encoding for HsGrpEL1_*WT*_ was transformed into the *E*.*coli* LOBSTR^55^ strain and grown in LB media containing 25*µ*g/mL kanamycin at 37°C until the OD600 reached 1.0. Protein expression was induced by addition of 200*µ*M IPTG and allowed to express for 3 hrs at 25°C. Cells were centrifuged, collected, and resuspended in buffer A with DNase I and RNAse I (Millapore Sigma) added prior to sonication. Cells were lysed using a Branson SFX550 sonifier on ice. The supernatant was loaded onto 5mL of Ni Sepharose High Performance resin (Cytiva) that had been pre-equilibrated with buffer A (without ZnCl_2_) supplemented with 10mM imidazole. The column was subsequently washed with buffer B containing 50mM imidazole until protein was no longer observed as detected by Bradford reagent. The target protein was eluted using 5 CVs of buffer B containing 300mM imidazole. Elutions were pooled and loaded onto 3mL of Strep-Tactin Superflow resin (IBA Lifesciences) that had been equilibrated with buffer E (20mM Tris pH 8, 150mM KCl, 5mM BME) and incubated for 30 min at 4°C. The column was washed with buffer E until no protein was observed in the eluent as detected by mixing with Bradford reagent. The target protein was eluted by application of 5 CVs of buffer E containing 2.5mM *d* -desthiobiotin (Sigma-Aldrich). Elutions were pooled, concentrated, and applied to a Superdex 200 Increase 10/300 GL (Cytiva) that had been pre-equilibrated with buffer D. Peak fractions were collected and analyzed using SDS-PAGE. The cleanest fractions were combined and concentrated to 0.8mg/mL. Concentrated fractions were used immediately, or flash frozen for storage.

#### GrpEL1_Y 173 A_

A pRSFDuet-1 vector encoding for the *Hs*GrpEL1_*Y* 173 *A*_ mutant was transformed into the *E*.*coli* LOBSTR^55^ strain. Recombinant protein expression and purification of GrpEL1_*Y* 173*A*_ was performed as described for GrpEL1_*WT*_.

Select peak fractions following SEC were visualized using SDS-PAGE. The cleanest fractions were combined and concentrated to 3.2 mg/mL. Concentrated fractions were used immediately, or flash frozen for storage.

#### Pam16/18 chimera

A pCDFDuet-1 vector encoding for the *Hs*Pam16/Pam18 chimera was transformed into *E*.*coli* LOBSTR cells and grown in LB media containing 25 *µ*g/mL of chloramphenicol and 50 *µ*g/mL of spectinomycin at 37°C until the OD600 reached 0.6. Protein expression was induced by addition of 100*µ*M IPTG and allowed to express for 16 hrs at 16°C. Cells were centrifuged, collected, and resuspended in buffer A containing DNase I and RNAse I (Millapore Sigma). Cells were lysed using a Branson SFX550 sonifier on ice. Lysed cells were centrifuged at 20,000 x g for 30 min at 4°C and the clarified supernatant was loaded onto 5mL of Ni Sepharose High Performance resin (Cytiva) that had been pre-equilibrated with buffer A that lacked ZnCl_2_. The column was washed with buffer B until protein was no longer observed in the eluent as detected by mixing with Bradford reagent. The target protein was eluted using 5 CVs of buffer B containing 300mM imidazole. Elutions were pooled and loaded onto 3mL of Amylose Resin (New England Biolabs) that had been equilibrated with buffer E and incubated for 1 hr at 4°C. The column was washed with buffer E until no protein was observed in the eluent as detected by Bradford reagent. The target protein was eluted by addition of 5 CVs of buffer E containing 10mM maltose. Elutions were pooled, concentrated, and applied to a Superdex 200 Increase 10/300 GL (Cytiva) that had been equilibrated with buffer D. Peak fractions were collected and analyzed using SDS-PAGE. The cleanest fractions were combined and concentrated to 1.0mg/mL. Concentrated fractions were used immediately, or flash frozen for storage.

### CryoEM sample preparation, data collection, and processing of the mortalin-GrpEL1_*W T*_ complex

Purified mortalin (20*µ*M), GrpEL1_*WT*_ (40*µ*M), and Pam16/Pam18 chimera (12*µ*M) were incubated at room temperature for 1hr with occasional mixing. The mixture was applied to a Superdex 200 Increase 10/300 GL (Cytiva) that was equilibrated with a buffer D. Fractions consisting of mortalin and GrpEL1_*WT*_, as visualized by SDS-PAGE, were concentrated to ∼0.3mg/mL. 4*µ*L of mortalin–GrpEL1_*WT*_ was applied to an UltraAuFoil 1.2/1.3, 300 grid that had been plasma-cleaned using a Gatan Solarus II plasma cleaner (10s, 15 Watts, 75% Ar/25% O2 atmosphere). Grids were manually blotted for ∼6 to 7s using Whatman No.1 filter paper before vitrification in a 1:1 ethane:propane liquid mixture cooled by liquid N_2_. The apparatus used was a custom manual plunge freezer designed by the Herzik lab located in a humidified (*>*95% relative humidity) cold room maintained at 4 °C.

Data acquisition was carried out at UCSD’s CryoEM Facility on a Titan Krios G4 (Thermo Fisher Scientific) operating at 300 keV equipped with a Selectris-X energy filter. Images were collected at a magnification of 130,000x in EFTEM mode (0.935 Å calibrated pixel size) on a Falcon 4 direct electron detector using a 10-eV slit width and a cumulative electron exposure of ∼60 electrons/ Å ^2^. Data were collected automatically using EPU with aberration free image shifting using a defocus range of -1.0 – -2.5 *µ*m. Initially, 952 movies were pruned from 1266 movies using a CTF acceptance range of ∼3.3-7.1 Å in cryoSPARC Live (cryoSPARC v4.3.1). Patch-based motion correction and CTF estimation were performed in cryoSPARC Live. 255,550 particles were selected using Blob Picker and extracted to a box size of 64 pixels (3.74 Å /pixel). Particles were subjected to two rounds of two-dimensional (2-D) classification (Round 1: 100 classes, 150 Å mask diameter, 200 Å outer mask diameter; Round 2: 50 classes, initial uncertainty factor: 1, 130 Å mask diameter, 160 Å outer mask diameter). Particles resembling a mortalin–GrpEL1_*WT*_ complex were used to generate an *ab initio* volume (no. of ab-initio classes: 2, maximum resolution: 6 Å, initial resolution: 15 Å, compute per-image optimal scales: on). This model revealed features of an intact mortalin–GrpEL1_*WT*_ complex, which was then used to create templates for template picking. 964,399 particles were identified using template-based particle picking and subjected to three sequential rounds of 2-D classification (50 classes, initial uncertainty factor: 1, 130 Å mask diameter, 160 Å outer mask diameter, no. of final full iterations: 2, batchsize per class: 1000). Classes containing false images were removed after iteration 1 while the best-behaving classes were retained in subsequent classifications.

In order to increase particle number and improve views, an additional 1757 movies were collected from the same batch of grids and subjected to the same patch-based motion correction and CTF estimation in cryoSPARC Live (as described above). 980,189 particles were selected using the previously generated templates and subjected to two sequential rounds of 2-D classification (100 classes, initial uncertainty factor: 1, 120 Å mask diameter, 150 Å outer mask diameter, batchsize per class: 1000). 703,658 particles were selected and combined with the previous dataset for a total of ∼1.35 million particles. These particles were sorted using five sequential rounds of 2-D classification (50 classes, 120 Å mask diameter, 150 Å outer mask diameter, number of final full iterations: 2, batchsize per class: 1000). Heterogeneous refinement, using an *ab initio* volume resembling the mortalin–GrpEL1_*WT*_ complex and a 20S proteasome volume (EMDB-8741), was performed. Particles locating to the mortalin–GrpEL1_*WT*_ complex class were re-extracted to a box size of 128 pixels (1.87 Å /pixel) and subjected to two additional rounds of heterogeneous refinement. This first round utilized a mortalin–GrpEL1_*WT*_ volume containing density for the SBD, a mortalin–GrpEL1_*WT*_ volume without density for the SBD, and a 20S proteasome volume (EMDB-8741) (force hard classification: on). The second round utilized the mortalin–GrpEL1_*WT*_ volumes, one containing and one lacking density for the SBD (force hard classification: on). 159,816 particles localizing to the class containing density for the SBD were re-extracted to a box size of 256 pixels (0.935 Å /pixel). Heterogeneous refinement, using the SBD-containing and SBD-lacking density mortalin–GrpEL14-_*WT*_ volumes, was performed and 132,552 particles classified into the SBD-containing class were selected. A soft mask of the SBD was generated (dilation radius: 7 pixels, soft padding width: 7 pixels) and used in 3-D classification (5 classes, force hard classification: on). Four classes (totaling 121,932 particles) were selected and subjected to heterogeneous refinement using two mortalin–GrpEL1_*W T*_ volumes, one with strong SBD density and one with weak SBD density. 66,065 particles classified into the strong SBD density class were re-extracted to a box size of 384 pixels (0.935 Å /pixel). Reference-based motion correction was performed on these particles using an extensive search for hyperparameters. Non-uniform refinement with CTF refinement turned on yielded a 3.06 Å resolution reconstruction.

To further increase particle number and improve views, an additional 1861 movies were collected as described above and subjected to patch-based motion correction and CTF estimation in cryoSPARC Live (as described above). Particles were identified using template-based particle picking and extracted to a 78 pixel box size (3.74 Å /pixel). Two sequential rounds of 2-D classification (50 classes, initial classification uncertainty factor: 1, 130Å mask diameter, 160 Å outer mask diameter, batchsize per class: 1000) were performed and 744,973 particles were selected. These particles were subjected to two rounds of heterogeneous refinement (default settings) using one class resembling the full mortalin–GrpEL1_*WT*_ complex and one junk class resembling noise. 248,553 particles classified into the mortalin–GrpEL1_*WT*_ class were reextracted to a box size of 256 pixels (0.935 Å /pixel). A 3 class *ab initio* reconstruction was performed on these particles resulting in 162,368 particles locating to a volume resembling the mortalin–GrpEL1_*WT*_ complex. Two rounds of heterogeneous refinement (Round 1: mortalin–GrpEL1_*WT*_ with strong SBD density, mortalin–GrpEL1_*WT*_ with weak SBD density; Round 2: mortalin–GrpEL1_*WT*_ with strong SBD density, noise density) were performed and 103,623 particles classified into the mortalin–GrpEL1_*WT*_ complex volume with strong SBD density were re-extracted to a box size of 384 pixels (0.935 Å /pixel). These particles were subjected to reference-based motion correction using an extensive search for hyperparameters. Particles from both reference-based motion correction jobs (56,380+96,395 particles) were combined and subjected to 3-D classification (4 classes). 150,347 particles locating to a volume resembling the mortalin–GrpEL1_*WT*_ complex were selected and subjected to 2-D classification (50 classes, initial uncertainty factor: 4, 130 Å mask diameter, 160 Å outer mask diameter, batch-size per class: 1000). 138,298 particles were selected and subjected to non-uniform refinement (window dataset (real-space): off, number of extra final passes: 2, initial lowpass resolution: 10 Å, minimize over per-particle scale: on, initialize noise model from images: on, dynamic mask near: 4 Å, dynamic mask far: 10 Å, optimize per-particle defocus: on, optimize per-group CTF params: on, fit spherical aberration: on, fit tetrafoil: on) resulting in a 2.96 Å resolution map.

### CryoEM sample preparation, data collection, and processing of the mortalin–GrpEL1_Y173 A_ complex

Purified mortalin (20*µ*M), GrpEL1_*Y* 173 *A*_ (40*µ*M), and Pam16/Pam18 chimera (12*µ*M) were incubated at room temperature for 1 hr with occasional mixing. The mixture was applied to a Superdex 200 Increase 10/300 GL (Cytiva) that was equilibrated with a buffer D. Fractions comprising mortalin and GrpEL1_*Y*173 *A*_, as visualized by SDS-PAGE, were concentrated to ∼0.5mg/mL. CryoEM grid preparation was prepared as described for the mortalin–GrpEL1_*WT*_ complex.

Data acquisition was carried out at UCSD’s CryoEM Facility on a Titan Krios G4 (Thermo Fisher Scientific) operating at 300 keV equipped with a Selectris-X energy filter. Images were collected at a magnification of 165,000x in EFTEM mode (0.735 Å calibrated pixel size) on a Falcon 4 direct electron detector using a 10-eV slit width and a cumulative electron exposure of ∼60 electrons/ Å^2^. Data were collected automatically using EPU with aberration free image shifting using a defocus range of -1.0 – -2.5 *µ*m. Patch-based motion cor-rection and CTF estimation were performed in cryoSPARC Live movies were filtered using a CTF acceptance range from 2-6 Å. 266,529 particles were selected using templates generated from the mortalin–GrpEL1_*WT*_ session, extracted to a box size of 96 pixels (3.74 Å /pixel), and subjected to three sequential rounds of 2-D classification (50 classes, initial uncertainty factor: 2, 130 Å mask diameter, 160 Å outer mask diameter, batchsize per class: 200). Three additional rounds of 2D classification were performed using the same parameters with the exception of an initial uncertainty factor of 4. 80,411 curated particles were then subjected to heterogeneous refinement using a mortalin–GrpEL1_*WT*_ *ab initio* class and a junk class. 41,043 particles localizing to the mortalin–GrpEL1_*WT*_ class underwent two sequential rounds of 2-D classification (25 classes, initial uncertainty factor: 4, 130 Å mask diameter, 160 Å outer mask diameter, number of final full iterations: 2, batchsize per class: 10000) and heterogeneous refinement using the mortalin–GrpEL1_*WT*_ *ab initio* class, a junk class, and a 20S proteasome class (EMDB-8741). 13,325 curated particles were then re-extracted to a box size of 384 pixels (0.735 Å /pixel).

An additional 1995 movies were collected from the same batch of grids and subjected to patch-based motion correction and CTF estimation using cryoSPARC Live. ∼1.25 million particles were identified using previously generated templates from mortalin–GrpEL1_*WT*_ and extracted to a box size of 96 pixels (3.74 Å /pixel). Six sequential rounds of 2-D classification (100 classes, initial uncertainty factor: 1, 130 Å mask diameter, 160 Å outer mask diameter, batchsize per class: 200) were performed, yielding 552,959 curated particles. These particles underwent heterogeneous refinement(force hard classification: on) using a mortalin–GrpEL1_*Y* 173 *A*_ volume, generated from the previous heterogeneous refinement, and a junk class. 252,261 particles localized to the mortalin–GrpEL1_*Y* 173 *A*_ volume and were subjected to 2-D classification (50 classes, initial uncertainty factor: 4, 130 Å mask diameter, 160 Å outer mask diameter), where 219,184 of the selected particles were subjected to another round of heterogeneous refinement using the same parameters described above. 150,060 particles assigned to the mortalin–GrpEL1_*Y* 173 *A*_ class underwent another round of 2-D classification (50 classes, initial uncertainty factor: 4, 130 Å mask diameter, 160 Å outer mask diameter). 130,080 particles were selected and utilized for a 3-class *ab initio* reconstruction that yielded 69,235 particles resembling mortalin–GrpEL1_*Y* 173 *A*_. These particles were re-extracted to a box size of 384 pixels (0.735Å/pixel). 13,325 and 69,235 unbinned particles from each dataset were combined (82,281) and subjected to non-uniform refinement (window dataset (real-space): off, number of extra final passes: 2, initial low-pass resolution: 10 Å, minimize over per-particle scale: on, initialize noise model from images: on, dynamic mask near: 4 Å, dynamic mask far: 10 Å, optimize per-particle defocus: on, optimize per-group CTF params: on, fit spherical aberration: on, fit tetrafoil: on, fit anisotropic mag.: on), yielding a 3.26 Å resolution EM map.

Our 3.26 Å cryoEM map exhibited anisotropy from dominant views and thus we collected an additional 1272 movies at a 10° tilt angle from the same batch of grids. Movies underwent patch-based motion correction and CTF estimation using cryoSPARC Live. Previously generated templates were used to pick 575,214 particles and were extracted to a box size of 96 pixels (3.74 Å /pixel). These particles were subjected to 2-D classification (100 classes, initial uncertainty factor: 1, 130Å mask diameter, 160Å outer mask diameter, batchsize per class: 200), resulting in 382,150 curated particles. These particles underwent heterogeneous refinement (force hard classification: on) using a mortalin–GrpEL1_*Y* 173 *A*_ volume and 20S proteasome class (EMDB-8741). 212,842 particles assigned to the mortalin–GrpEL1_*Y* 173 *A*_ complex class were subjected to an additional round of 2-D classification (50 classes, initial uncertainty factor: 2, 130Å mask diameter, 160Å outer mask diameter, batchsize per class: 200) where 165,148 selected particles subsequently underwent heterogeneous refinement (force hard classification: on) using a mortalin–GrpEL1_*Y* 173 *A*_ class, a 20S proteasome class (EMDB-8741), and a junk class. 87,397 particles localizing to the mortalin–GrpEL1_*Y* 173 *A*_ class underwent a 2-class *ab initio* reconstruction (max resolution: 12Å, initial resolution 35Å) where 51,461 particles locating to a volume resembling a mortalin–GrpEL1_*Y* 173 *A*_ complex were selected. These particles underwent heterogeneous refinement (force hard classification: on) using one mortalin–GrpEL1_*WT*_ class with strong SBD density and one mortalin–GrpEL1_*WT*_ class with weak SBD density. 38,765 particles localizing to the complex class with strong SBD density were then re-extracted to a box size of 384 pixels (0.735Å/pixel). A soft mask of the SBD was generated (dilation radius: 4.41 pixels, soft padding width: 6 pixels) and used in a focused 3-D classification (2 classes, target resolution: 6Å, initial structure low-pass resolution: 20Å) with combined particles from the three mortalin–GrpEL1_*Y* 173 *A*_ datasets (13,325+69,235+38,765). One volume appeared to contain mortalin–GrpEL1_*Y* 173 *A*_ with an intact SBD lid (mortalin–GrpEL1_*Y* 173*A*_) whereas the second volume contained mortalin–GrpEL1_*Y* 173 *A*_ without the SBD-lid (mortalin–GrpEL1_*Y* 173 *A*_ -lid). These two volumes, in addition to a junk volume, were used as references for a heterogeneous refinement (force hard classification: on) using the combined particles and yielded 51,324 particles in the mortalin–GrpEL1_*Y* 173 *A*_ class and 54,402 particles in the mortalin–GrpEL1_*Y* 173 *A*_ -lid class. Non-uniform refinement (window dataset (real-space): off, number of extra final passes: 2, initial lowpass resolution: 10Å, minimize over per-particle scale: on, initialize noise model from images: on, dynamic mask near: 4Å, dynamic mask far: 10Å, optimize per-particle defocus: on, optimize per-group CTF params: on, fit spherical aberration: on, fit tetrafoil: on, fit anisotropic mag.: on) was performed on each of these classes, yielding EM maps with nominal resolutions of 3.38Å.

### Model Building of mortalin-GrpEL1_*WT*_

To model the mortalin NBD and GrpEL1 regions of our mortalin–GrpEL1_*WT*_ complex, we initially referenced the crystal structure of *E. coli* DnaK-GrpE (PDB ID: 1DKG).^22^ For the mortalin IDL and SBD, we referenced the crystal structure of the *Hs*Hsp70-SBD bound to a peptide substrate (PDB ID: 4PO2). These models were used as initial structures and docked as rigid bodies into our 2.96Å EM map.

For mortalin we modeled residues 47-639. For GrpEL1-A, residues 64-217 were modeled in addition to a C-terminal serine and alanine representing the linker region. For GrpEL1-B, residues 62-212 were modeled. The bound mortalin substrate was modeled with residues 434-440 representing the interdomain linker. All models were subjected to manual curation and adjustment in COOT followed by real-space refinement in PHENIX (version 1.21). Model figures were generated in UCSF ChimeraX.

### Model Building of mortalin-GrpEL1_*Y* 173 *A*_

#### mortalin-GrpEL1_Y 173 A_

To model our mortalin-GrpEL1_*Y* 173 *A*_ structure, we performed rigid-body docking using our mortalin-GrpEL1_*WT*_ model on the mortalin NBD, IDL, GrpEL1-A and GrpEL1-B regions. The SBD and bound substrate regions were docked independently to accommodate the mortalin-GrpEL1_*Y* 173 *A*_ SBD EM density. All models were subjected to manual curation and adjustment in COOT followed by real-space refinement in PHENIX. Model figures were generated in UCSF ChimeraX.

#### mortalin-GrpEL1_Y 173 A_ -lid

For the mortalin-GrpEL1_*Y* 173 *A*_ -lid structure, the lower resolution EM density for mortalin’s SBD did not allow for de novo placement of all residues. Therefore, subdomains from the mortalin-GrpEL1_*Y* 173 *A*_ structure were rigid body docked and reference-based refinement parameters in PHENIX were used. All models were subjected to manual curation and adjustment in COOT followed by real-space refinement in PHENIX. Model figures were generated in UCSF ChimeraX.

### All-atom molecular dynamics (MD) simulations of the mortalin-GrpEL1_*WT*_ and mortalin-GrpEL1_*Y* 173 *A*_ complexes using NAMD

#### System Preparation

Coordinates for the mortalin-GrpEL1_*WT*_ and mortalin–GrpEL1_*Y* 173 *A*_ complexes were defined using the PDB coordinate files following refinement into the corresponding cryoEM maps. These models were solvated with explicit TIP3 water molecules and the number of Na^+^ and Cl^−^ ions were adjusted to neutralize the system charge (with an ionic strength set to 150mM). The total number of atoms for the final systems were 129,167 for mortalin-GrpEL1_*W T*_ and 137,771 for mortalin-GrpEL1_*Y* 173 *A*_ and orthorhombic periodic cells of 85 Å × 123 Å × 122 Å and 89 Å × 125 Å × 132 Å, respectively.

All-atom MD simulations were performed using the CUDA memory-optimized version of NAMD 2.14^56^ and CHARMM36m force fields.^57^ Each simulation began with an energy minimization step to remove any steric clashes and to ensure that the system was in a low-energy state before dynamic simulations. A minimization of 2000 steps was performed with the following key settings: a cutoff of 12.0Å for nonbonded interactions, a pair list distance of 14.0Å, and the use of the Particle Mesh Ewald (PME) method for long-range electrostatics with a grid spacing of 1Å.^58^ Switching was enabled with a switch distance of 10.0Å, and all atoms were wrapped to the primary simulation cell, including water molecules, to maintain periodic boundary conditions. Following minimization, the system underwent a temperature annealing process. The temperature was gradually increased from 60K to 300K to allow the system to adapt slowly to the target temperature, reducing the potential for introducing artifacts due to rapid temperature changes. This process was managed through Langevin dynamics with a damping coefficient of 1 ps^−1^, ensuring temperature and pressure stabilization throughout the annealing phase. The system was equilibrated at 300K for a duration sufficient to ensure that all parts of the system had reached a stable state **(Supplementary Figure 18A)**. The equilibration phase used the same cutoff, pair list distance, and PME settings as the previous steps to maintain consistency in the treatment of non-bonded interactions. Langevin dynamics continued to control the temperature, with a target pressure of 1.01325 bar maintained via a Langevin piston. The simulation time for this phase was extended to 500,000 steps for both structures to ensure comprehensive equilibration. Harmonic constraints were applied during minimization, annealing, and equilibration to maintain the structural integrity of the starting complexes.

Following minimization, heating, and equilibration, the systems were submitted to productive MD simulations under NPT conditions. A timestep of 2 fs was used, with non-bonded interactions (van der Waals and short-range electrostatic) calculated at each timestep using a cutoff of 12 Å and a switching distance of 10 Å. All simulations employed periodic boundary conditions, utilizing the particle-mesh Ewald method with a grid spacing of 1 Å to evaluate long-range electrostatic interactions. The MD production was carried out for 75,000,000 steps, equivalent to 150 ns, with coordinates saved every 1000 steps (2 ps). Randomized triplicates were run for each structure; 0.45 *µ*s (75,000 frames) combined total for each structure.

#### RMSD Calculation

The root mean square deviation (RMSD) of the mortalin-GrpEL1_*WT*_ and mortalin–GrpEL1_*Y* 173 *A*_ complexes was calculated to assess the structural stability and conformational changes of the proteins during the simulations. The RMSD analysis was performed using the Visual Molecular Dynamics (VMD) software.^59^ The initial structure of the protein was used as the reference structure for the RMSD calculations. The RMSD values were computed for the backbone atoms (N, C α, C, and O, not including hydrogens) of the protein after aligning the structures to the reference frame to remove translational and rotational movements. The RMSD analysis was carried out for the entire trajectory, and the results are plotted in **Supplementary Figure 18B** to visualize the structural fluctuations and stability of each complex.

#### Angular Distribution Calculation

To investigate the structural dynamics of mortalin-GrpEL1_*WT*_ and mortalin-GrpEL1_*Y* 173 *A*_ during the molecular dynamics simulations, we calculated the angular motion between two predefined vectors for each simulation frame. For mortalin-GrpEL1_*WT*_, vector 1 (v1) extended from the C α atom of residue 570 in the SBD α lid to the C α atom of residue 596 within the same domain, while Vector 2 (v2) spanned from the C α atom of residue 570 in the SBD α lid to the C α atom of residue 101 in the GrpEL1-B long α -helix. These vectors were chosen to monitor the lateral movement of the SBD lid relative to the wing domain.

For mortalin-GrpEL1_*Y* 173 *A*_, vector 1 (v1) was defined from the C α atom of residue 562 to the C α atom of residue 511, and vector 2 (v2) connected the C α atom of residue 562 in the SBD to the C α atom of residue 596 in the SBD lid. These vectors tracked the medial movement of the SBD lid with respect to the SBD.

The angle between v1 and v2 was determined at each timestep using a custom Tcl script in the Visual Molecular Dynamics (VMD) software. The script iteratively updated the coordinates of the selected C*α* atoms, calculated the angle between the two vectors, and recorded the angle along with the corresponding frame number. Vectors are visualized in **Supplementary Figure 13**.

The angular data was then visualized using MATLAB^60^ to depict the trajectory of these angles throughout the simulation. The angles were plotted against the step number, with kernel density estimation applied to the data to generate a smooth distribution curve.

### Elastic Network Model Analysis Using ProDy

#### System Preparation and Model Construction

The mortalin-GrpEL1_*WT*_ and the mortalin-GrpEL1_*Y* 173 *A*_ structures were parsed using ProDy,^61^ with secondary structures assigned automatically. Alpha carbon (C α) selections were made using all C α in both conformations.

#### Anisotropic Network Model (ANM) Calculation

ANM was used for both conformations to explore the protein’s dynamics. The Hessian matrix was constructed to capture the 3D anisotropic behaviors of the protein’s movements.

A cutoff distance of 10 Å for interactions, a gamma value of 1 indicating the strength of spring constants in the model. The number of modes calculated was set to 10 to focus on the most significant low-frequency motions that correspond to functional movements within the protein. The analysis was conducted with backbone C α atoms to emphasize the core structural dynamics.

#### Mode Analysis and Visualization

The ANM modes were analyzed through visualization in VMD using the Normal Mode Wizard (NMWiz) plugin. The ANM model, initially based on C α atoms, was extended to all atoms to refine the analysis, and include more detailed atomic interactions. This extension was performed using ProDy’s^61^ extend mode method, which bridges the coarse-grained ANM representation with a detailed all-atom model. This facilitated an intuitive understanding of the modes’ physical implications on the protein’s structure and function.

All 10 modes were visualized and analyzed for insights into the protein’s intrinsic motions. Mean square fluctuations and cross-correlations based on the computed modes were also examined to understand the distribution of movements and the relationship between different regions of the protein.

## Data Availability

The data supporting this study are available from the corresponding author upon request. All models and associated cryoEM maps have been deposited into the Electron Microscopy Data Bank (EMDB) and the PDB. The depositions include final maps, unsharpened maps, half maps, and associated FSC curves. The accession codes are listed here and in Table 1. Mortalin-GrpEL1_*WT*_ : EMD-44675 and PDB: 9BLS; Mortalin-GrpEL1_*Y* 173 *A*_: EMD-44676 and PDB: 9BLT; Mortalin-GrpEL1_*Y* 173 *A*_ -lid: EMD-44677 and PDB: 9BLU. The all-atom molecular dynamics simulations system files and run time files are available on Zenodo under accession number XXXXXX.

## Ethics Declaration

The authors declare no competing interests.

## Acknowledgements

We are grateful to Prof. Rommie Amaro and Prof. Kevin Corbett for providing valuable feedback on this manuscript, and the entirety of the Herzik lab for facilitating insightful discussions. We also thank Dr. Brian Cook for his generous mentorship on protein purification, cryoEM sample preparation, cryoEM data collection, cryoEM data processing, and preparation of the manuscript. We are grateful to Sam Marchant for producing the R126W point mutation of mortalin. We also thank members of the University of California, San Diego (UCSD)’s Cryo-EM Facility and the broader cryoEM community at UCSD, and Brendan Dennis, Kevin Smith, and the UCSD Physics Computing Facility for their insights and support. Molecular graphics and analyses were performed with UCSF ChimeraX, developed by the Resource for Biocomputing, Visualization, and Informatics at the University of California, San Francisco (UCSF), with support from National Institutes of Health (NIH) grant R01-GM129325 and the Office of Cyber Infrastructure and Computational Biology, National Institute of Allergy and Infectious Diseases.

