## Supplementary Information for "Structural insights into GrpEL1-mediated nucleotide and substrate release of human mitochondrial Hsp70"

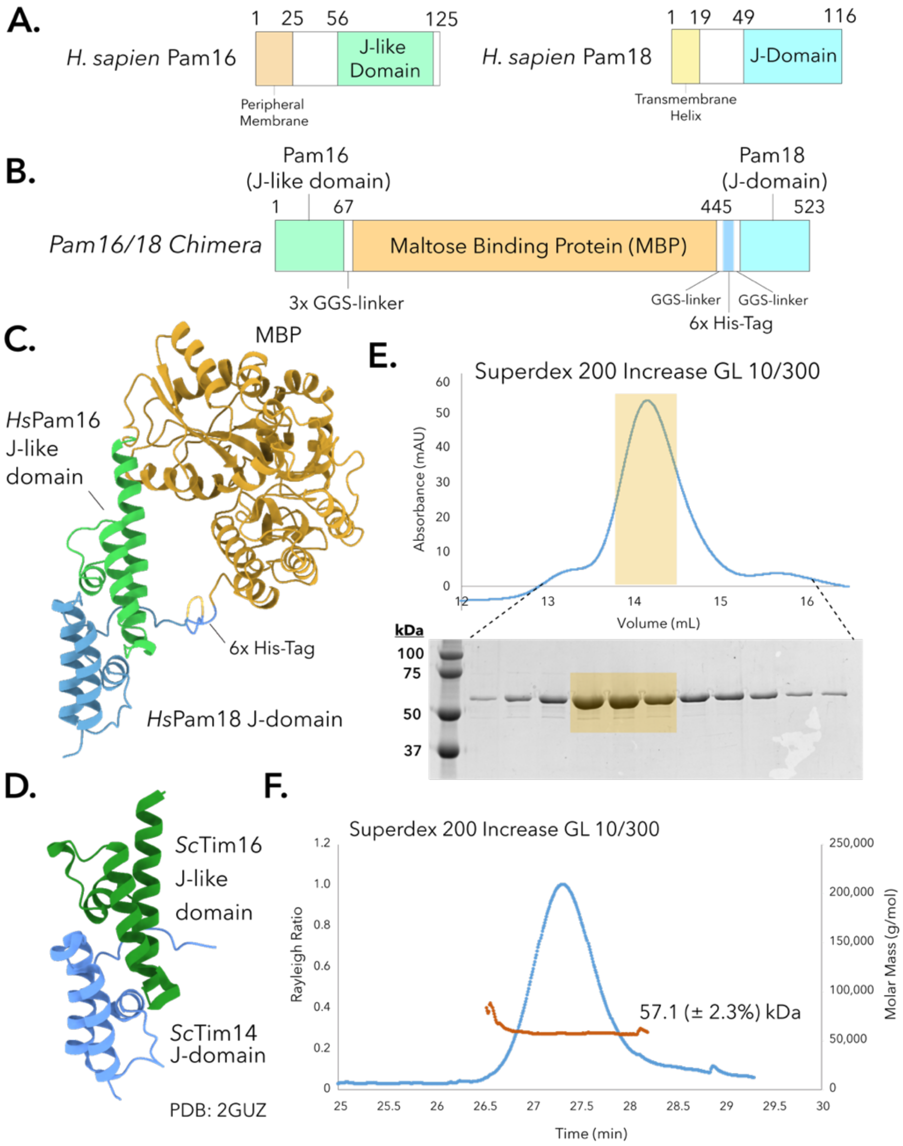


**Supplementary Figure 1. Rational construct design and biochemical preparation of a chimeric human Pam16/Pam18 fusion. A.** Domain topology of *Hs*Pam16 and *Hs*Pam18. **B.** Domain topology of the designed Pam16/Pam18 chimera fusion. **C.** AlphaFold2 prediction of the Pam16/Pam18 chimera shown in **B**. **D.** Crystal structure of the *Saccharomyces cerevisiae* Tim16 J-like domain (*Hs*Pam16 homolog) and Tim14 J-domain (*Hs*Pam18 homolog) (PDB: 2GUZ). The arrangement of the Tim16/Tim14 heterodimer is similar to the predicted structure of the Pam18/Pam16 chimera. **E.** Preparatory size exclusion chromatogram of the Pam16/Pam18 chimera and SDS-PAGE analysis. The highlighted regions were pooled and concentrated for immediate use or flash frozen using liquid nitrogen. **F.** Size exclusion chromatography multi-angular light scattering (SEC-MALS) analysis of purified Pam16/Pam18 chimera. Expected molecular weight: 57 kDa. The light scattering curve is represented in blue and the molecular weight determination is represented in red (with estimated error shown in parentheses).

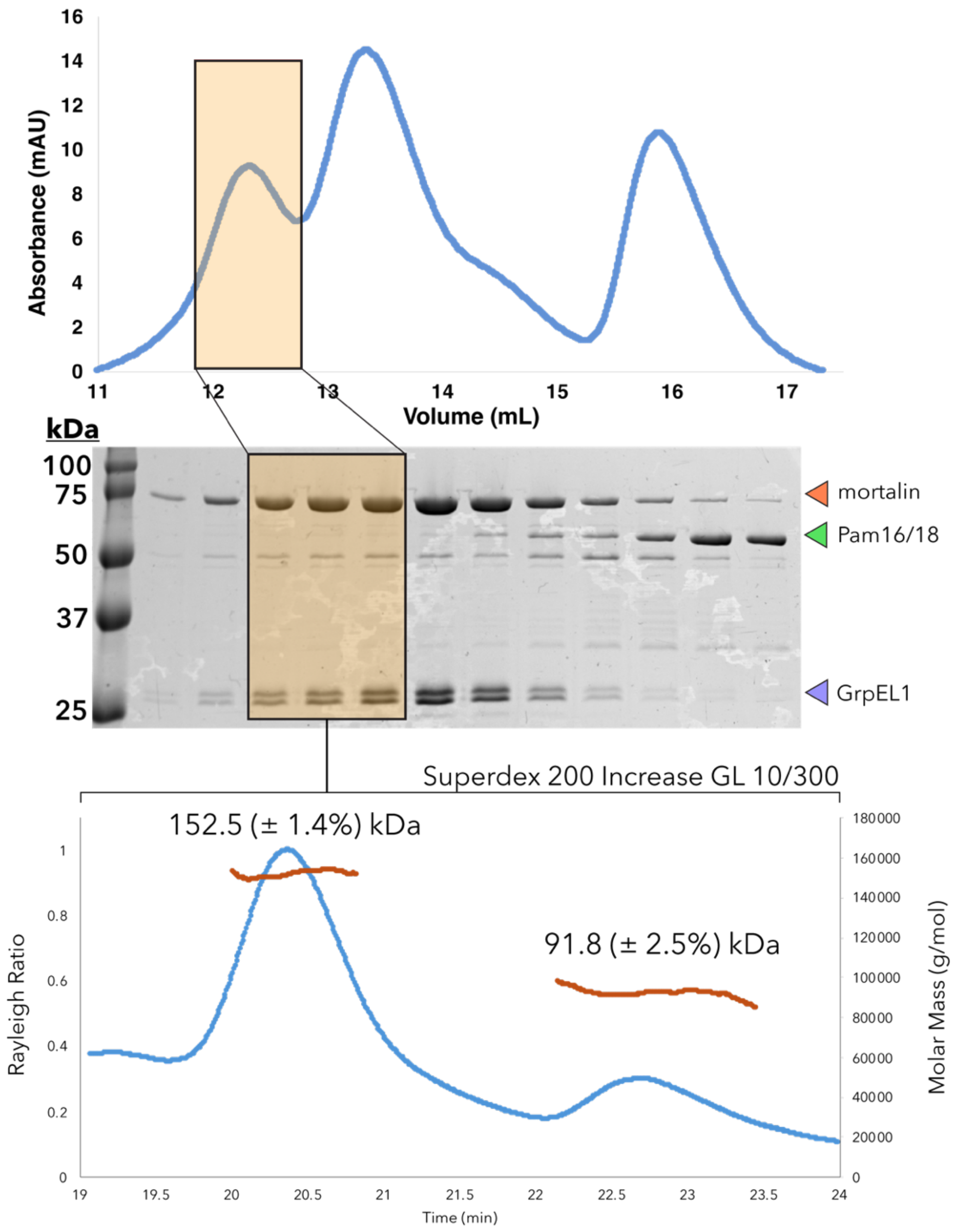


**Supplementary Figure 2. Biochemical preparation of the mortalin-GrpEL1$_{WT}$ complex.** *(top)* Highlighted fractions from a Superdex 200 Increase GL 10/300 SEC run (adapted from Figure 1B) were concentrated to 0.9 mg/mL and subsequently subjected to *(bottom)* SEC-MALS analysis (Superdex 200 Increase GL 10/300 at 0.5 mL/min at room temperature). The higher molecular weight species corresponds to full-length mortalin (70.2 kDa), a GrpEL1 homodimer (47 kDa), and a mortalin IDL-SBD truncation product (28.5 kDa). The lower molecular weight species corresponds to a mortalin NBD (41.8 kDa) and a GrpEL1 dimer (47 kDa). Mortalin: 70.2 kDa, GrpEL1 (monomer): 23.5 kDa, mortalin (NBD): 41.8 kDa, mortalin (IDL-SBD): 28.5 kDa. The light scattering curve is represented in blue and the molecular weight determination is represented in red (with estimated error shown in parentheses).

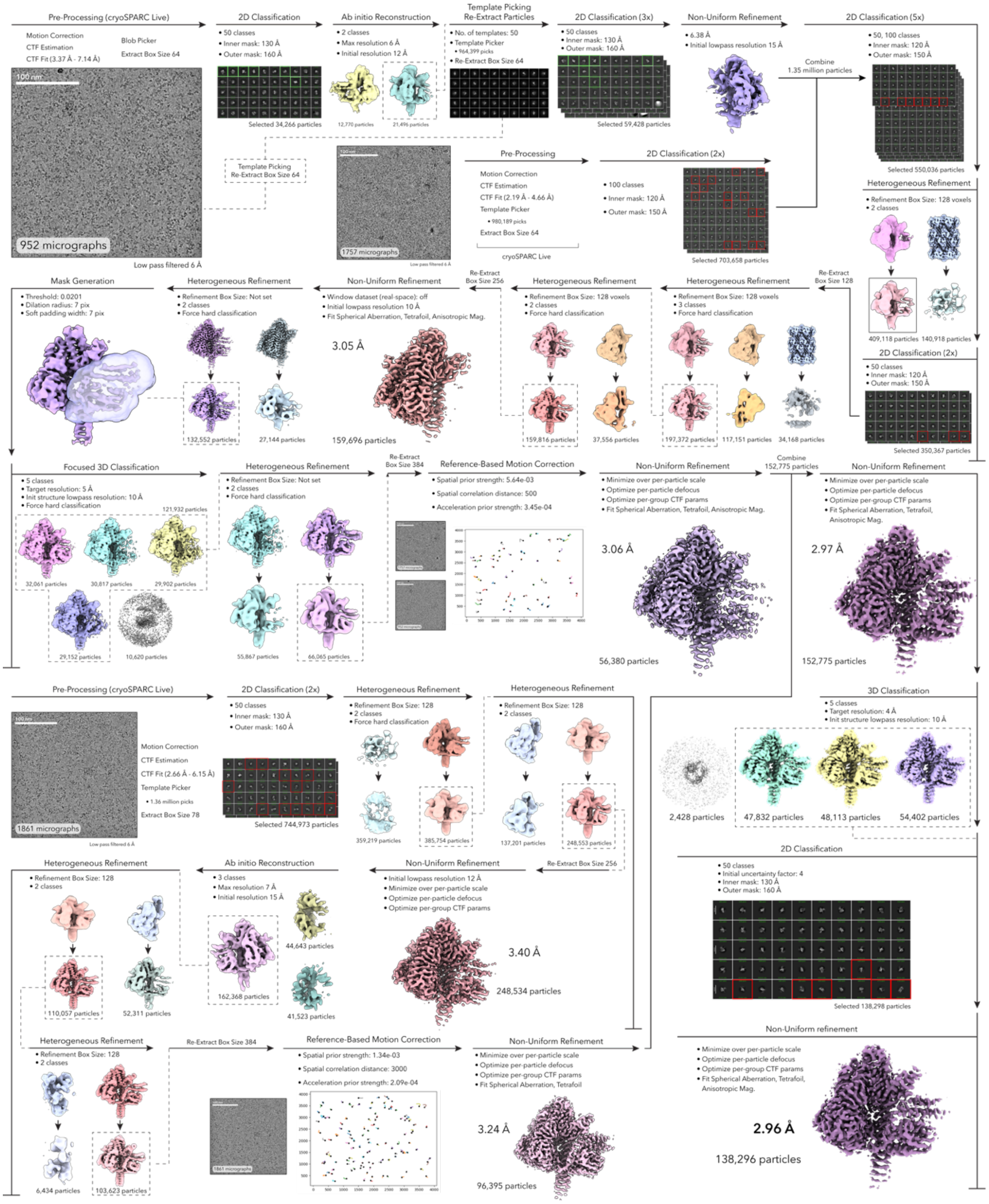


**Supplementary Figure 3. CryoEM data processing of the mortalin-GrpEL1$_{WT}$ complex.** See ***Methods*** for detailed explanation of cryoEM data processing. For 2-D classification, classes highlighted in green were chosen as selected particles; classes highlighted in red were excluded from the selected particles. Relevant parameters for each step are listed accordingly.

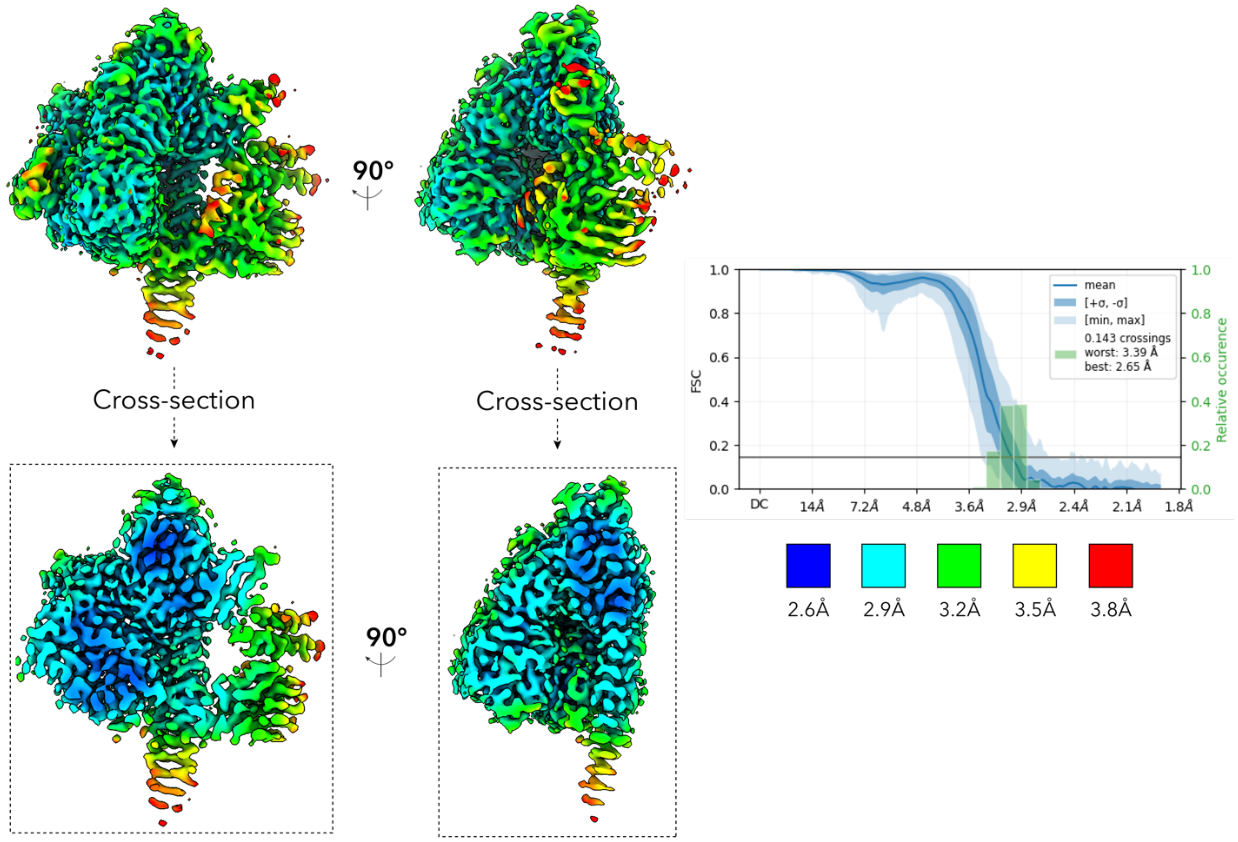


**Supplementary Figure 4. Local resolution estimation of the mortalin-GrpEL1$_{WT}$ complex.** Locally filtered EM density of the mortalin-GrpEL1$_{WT}$ complex colored by local resolution. Cross-sections of the EM density are shown below each view. 3-D Fourier shell correlation (FSC) plots generated from the independent half maps contributing to the ~2.96 Å mortalin-GrpEL1$_{WT}$ map.

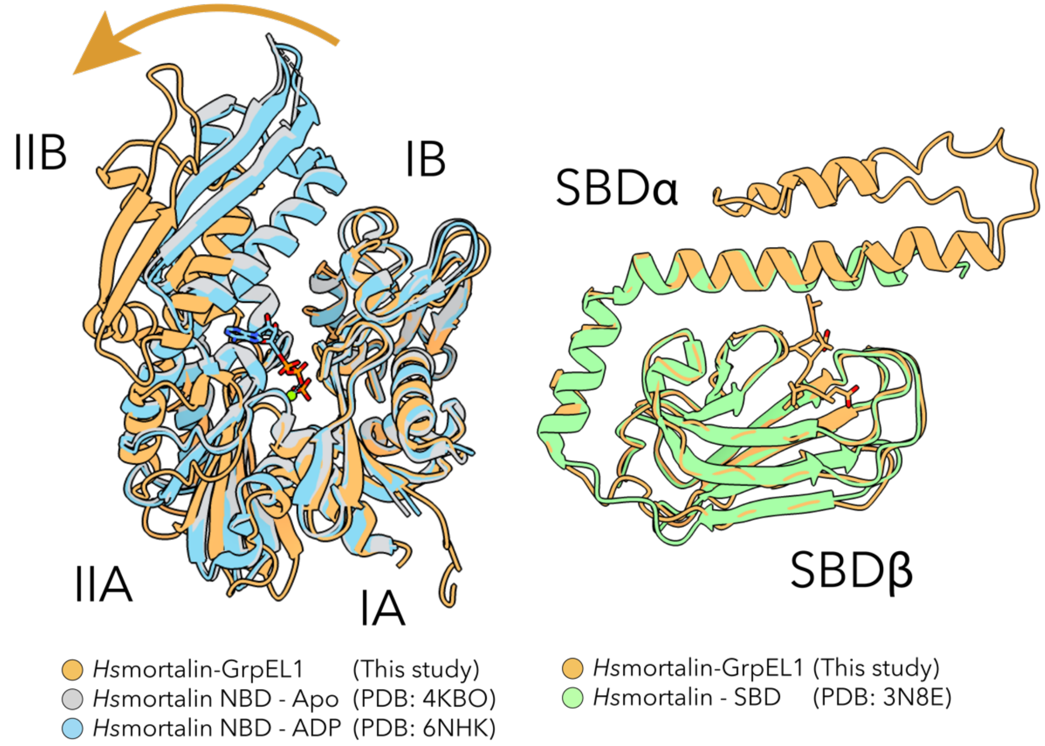


**Supplementary Figure 5. Comparison of the NBD and SBD domains in the mortalin-GrpEL1$_{WT}$ complex with mortalin crystal structures.** *(left)* Superposition of the NBD from *Hs*mortalin complexed with GrpEL1$_{WT}$ (this study) and *Hs*mortalin NBD crystal structures in either apo (PDB: 4KBO) or ADP-bound (PDB: 6NHK) states. *(right)* Superposition of the substrate-bound SBD from *Hs*mortalin complexed with GrpEL1$_{WT}$ (this study) and the *Hs*mortalin SBD crystal structure (PDB: 3N8E).

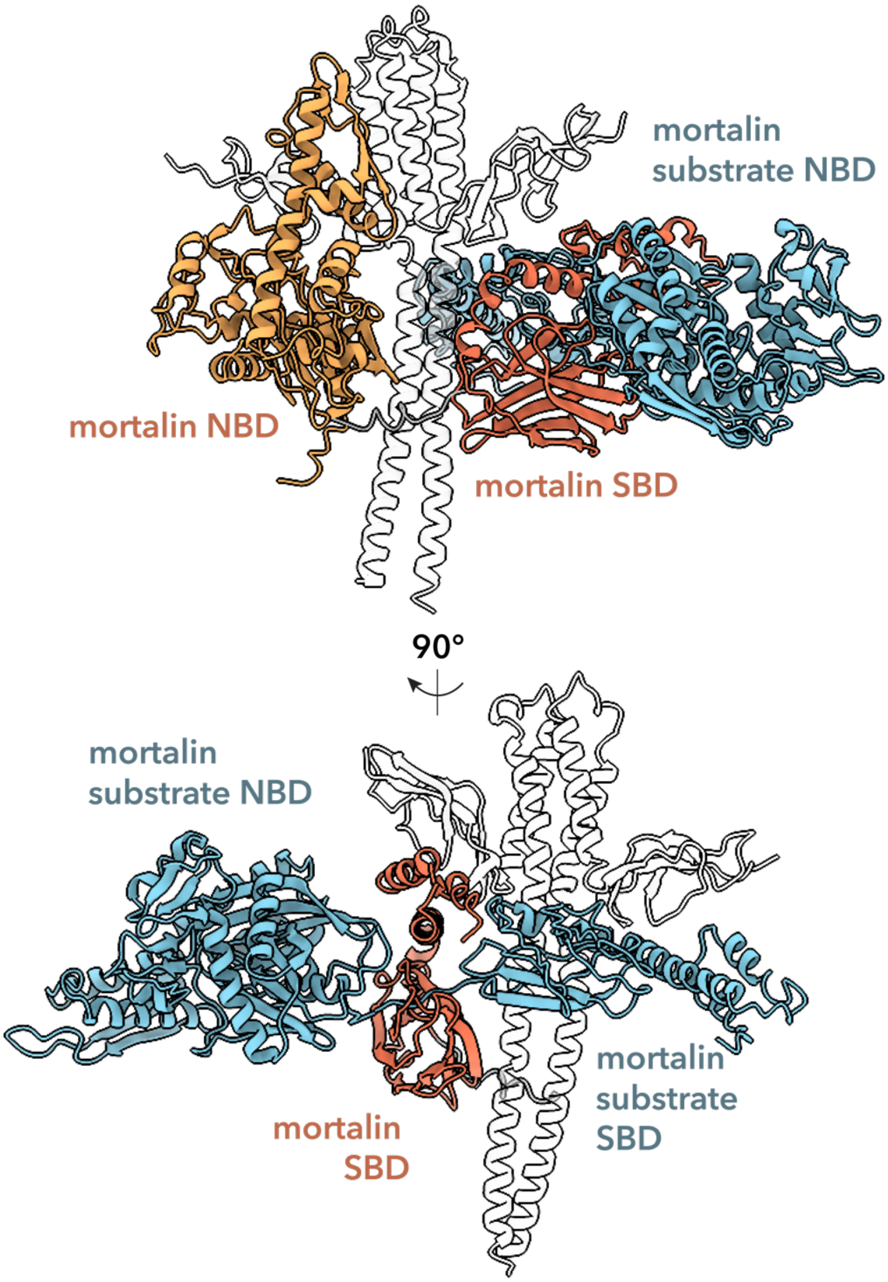


**Supplementary Figure 6. Mortalin in complex with GrpEL1 can accommodate full-length mortalin as a substrate.** The mortalin SBD was rigid-body docked into the posterior substrate EM density with a low-pass filtered mortalin-GrpEL1$_{WT}$ map, as shown in **Figure 4B**. The NBD from the mortalin-GrpEL1$_{WT}$ structure was separated and aligned with the IDL in the mortalin-GrpEL1$_{WT}$ complex structure. The NBD was oriented to mimic substrate binding of a full-length mortalin unit within the mortalin SBD of the mortalin-GrpEL1$_{WT}$ complex structure. An orthogonal view shows that the complex mortalin SBD could accommodate full-length mortalin.

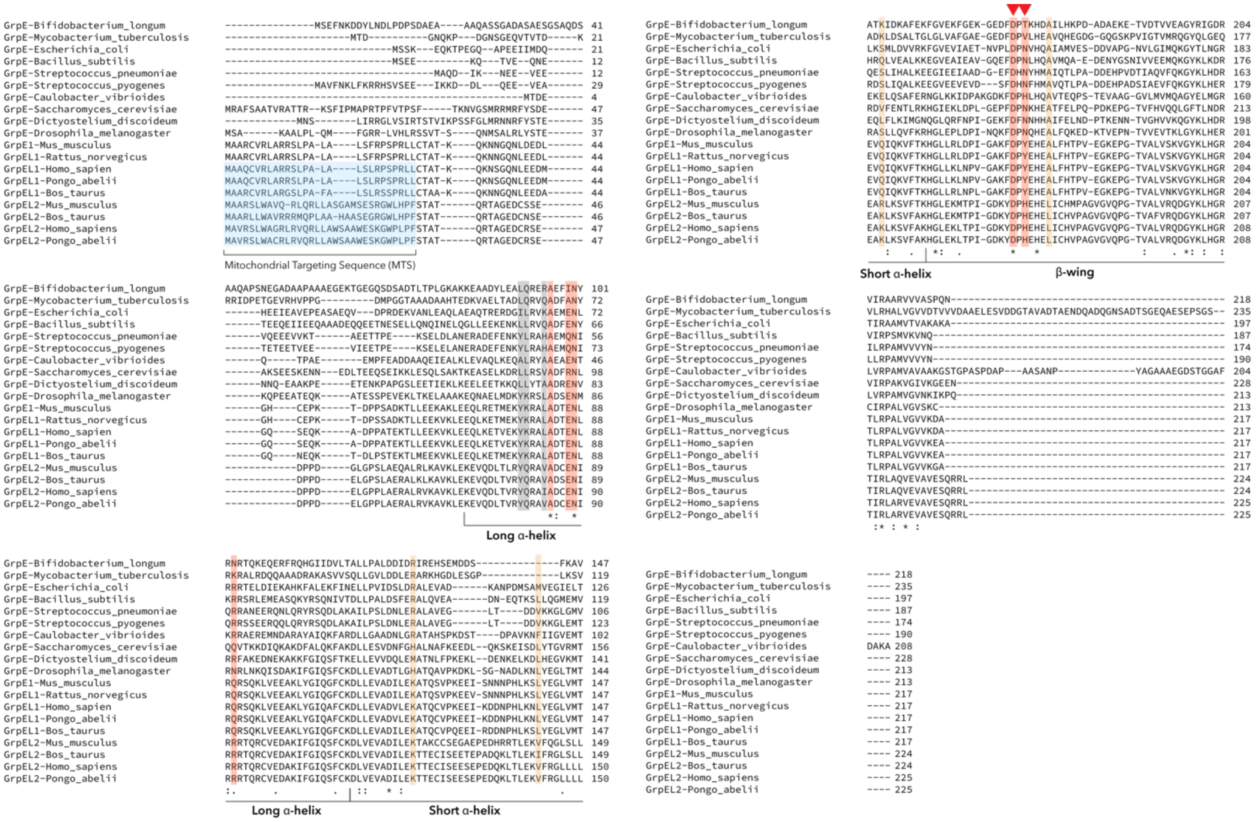

**Supplementary Figure 7. Multiple sequence alignment of GrpE-like homologs.** Multiple sequence alignment of 19 GrpE-like homologs. Residues that interact with the mortalin NBD are colored in light orange, residues that interact with the mortalin IDL are colored in grey, residues that interact with the SBD are colored in red. D171 and Y173 that interact with the mortalin SBDα lid are indicated with a red triangle. The mitochondrial targeting sequence (MTS) is colored in light blue in mitochondrially targeted GrpE-like species. Residues that are fully conserved across species are noted with an asterisk (*). Residues with strong conservation of similar properties are noted with a colon (:). Residues with weak conservation of similar properties are noted with a period (.). Multiple sequence aligned was performed in Clustal Omega. *Bifidobacterium longum* (entry: Q8G6W2), *Mycobacterium tuberculosis* (entry: P9WMT5), *Escherichia coli* (entry: P09372), *Bacillus subtilis* (entry: P15874), *Streptococcus pneumoniae* (entry: Q97S73), *Streptococcus pyogenes* (entry: Q5XAD5), *Caulobacter vibrio* (entry: B8GXP4), *Saccharomyces cerevisiae* (entry: P38523), *Dictyostelium discoideum* (entry: Q54QF9), *Drosophila melanogaster* (entry: P48604), *Mus musculus* (GrpEL1) (entry: Q99LP6), *Rattus norvegicus* (entry: P97576), *Homo sapien* (GrpEL1) (entry: Q9HAV7), *Pongo abelii* (GrpEL1) (entry: Q5RA81), *Bos taurus* (GrpEL1) (entry: Q3SZC1), *Mus musculus* (GrpEL2) (entry: O88396), *Bos taurus* (GrpEL2) (entry: Q0P5N5), *Homo sapien* (GrpEL2) (entry: Q8TAA5), *Pongo abelii* (GrpEL2) (entry: Q5R435).

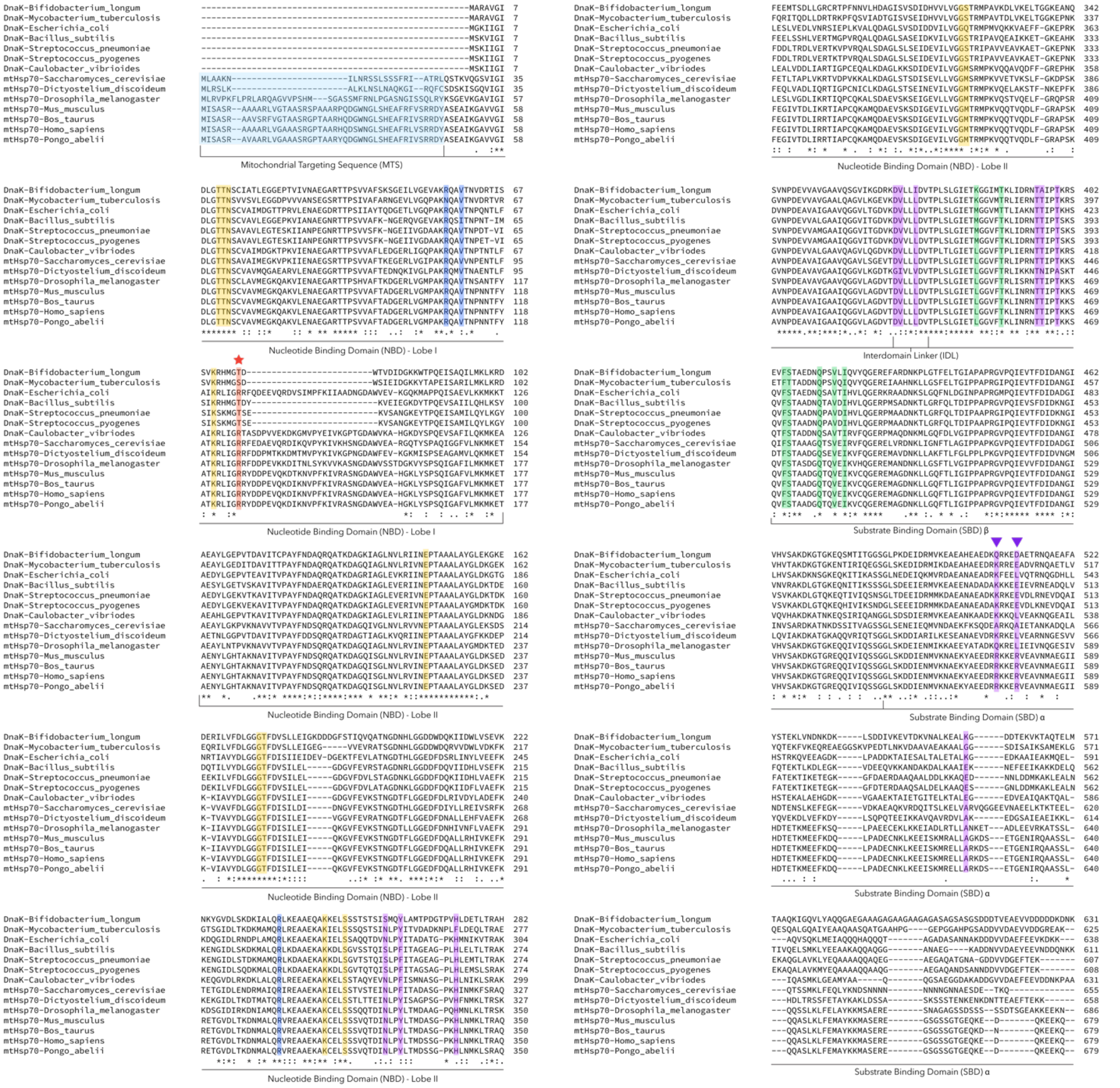


**Supplementary Figure 8. Multiple sequence alignment of mitochondrial (mt)Hsp70 homologs.** Multiple sequence alignment of 14 mtHsp70 homologs. Residues that interact with GrpEL1-A are colored in dark blue and residues that interact with GrpEL1-B are colored in purple. R574 and R578 that interact with GrpEL1-B are indicated with a purple triangle. R126, mutated to R126W in this study, is highlighted in orange and indicated with an orange star. Residues that interact with bound substrate are colored in green. Residues in that nucleotide binding site are colored in gold. The mitochondrial targeting sequence (MTS) is colored in light blue in mitochondrially targeted mtHsp70 species. Residues that are fully conserved across species are noted with an asterisk (*). Residues with strong conservation of similar properties are noted with a colon (:). Residues with weak conservation of similar properties are noted with a period (.). Multiple sequence alignment was performed in Clustal Omega. *Bifidobacterium longum* (entry: B7GT47), *Mycobacterium tuberculosis* (entry: P9WMJ9), *Escherichia coli* (entry: P0A6Y8), *Bacillus subtilis* (entry: P17820), *Streptococcus pneumoniae* (entry: Q8CWT3), *Streptococcus pyogenes* (entry: P0C0C6), *Caulobacter vibrioides* (entry: P20442), *Saccharomyces cerevisiae*

(entry: P0CS90), *Dictyostelium discoideum* (entry: Q8I0H7), *Drosophila melanogaster* (entry: P82910), *Mus musculus* (entry: P38647), *Bos taurus* (entry: Q3ZCH0), *Homo sapien* (entry: P38646), *Pongo abelii* (entry: Q5R511).

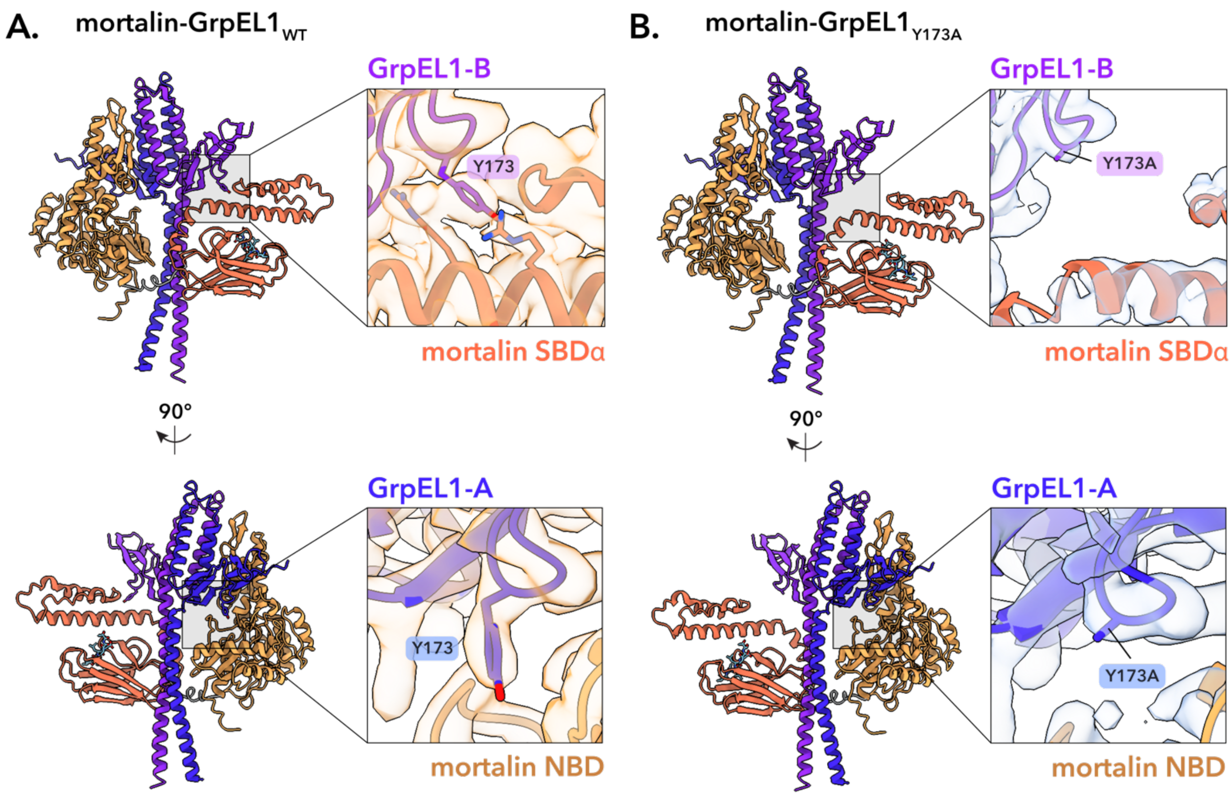


**Supplementary Figure 9. GrpEL1 exhibits interaction interfaces that uniquely interact with either the mortalin NBD or SBD. A.** Visualization of Y173 in both GrpEL1-B *(top)* and GrpEL1-A *(bottom)* in the mortalin-GrpEL1$_{WT}$ structure. While Y173 makes close contacts with mortalin in GrpEL1-B, Y173 does not appear to contact mortalin in GrpEL1-A. The mortalin-GrpEL1 map was modified using DeepEMhancer for visualization. **B)** Visualization of Y173A in both GrpEL1-B *(top)* and GrpEL1-A *(bottom)* in the mortalin-GrpEL1$_{Y173A}$ structure. Large separation is observed between Y173A in GrpEL1-B and mortalin. Y173A in GrpEL1-A does not appear to contact mortalin. The mortalin-GrpEL1$_{Y173A}$ map was modified using DeepEMhancer for visualization.

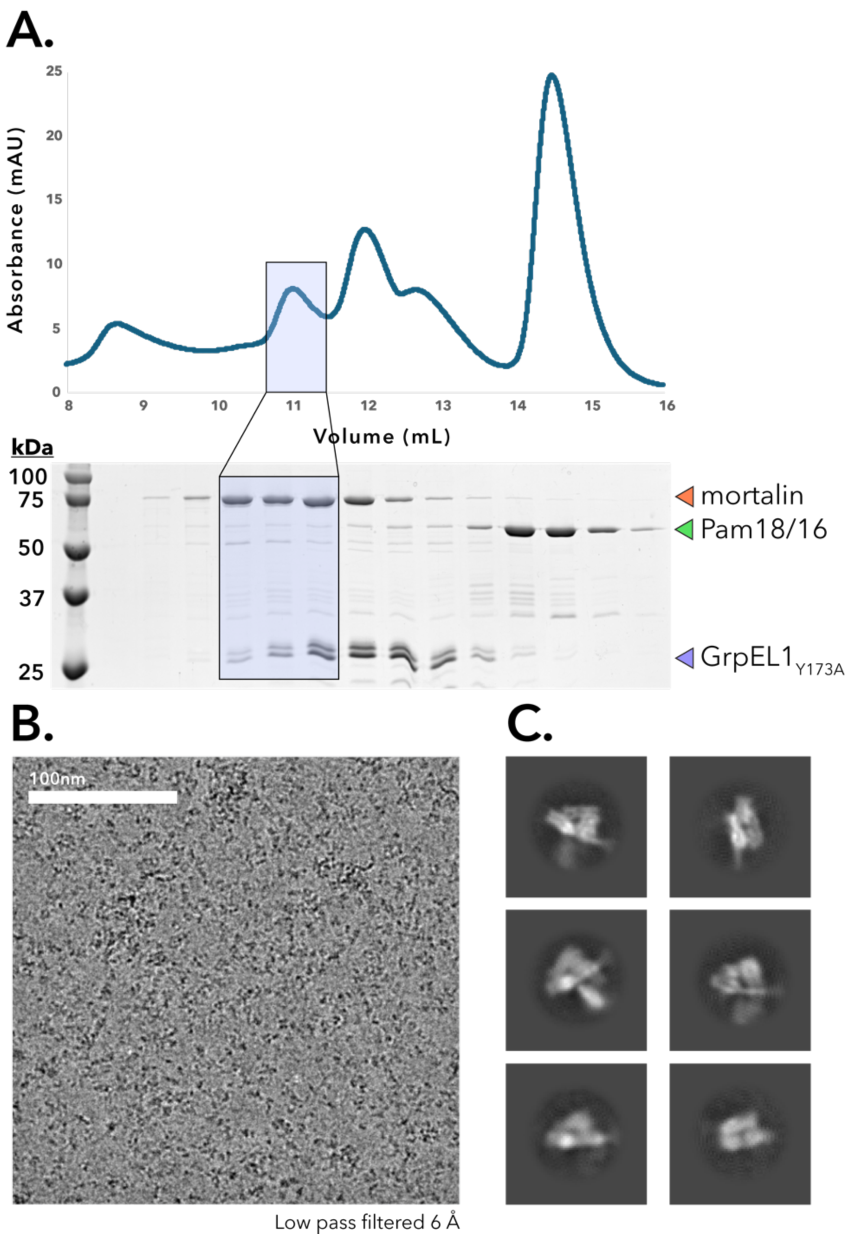


**Supplementary Figure 10. Biochemical preparation of the mortalin-GrpEL1$_{Y173A}$ complex. A.** Size exclusion chromatogram ($A_{280}$) and corresponding SDS-PAGE analysis. The left-most peak was concentrated and vitrified onto cryoEM grids. **B.** Representative micrograph of the mortalin-GrpEL1$_{Y173A}$ complex. Micrograph was low-pass filtered to 6 Å for clarity. **C.** Representative two-dimensional (2-D) classes of the mortalin-GrpEL1$_{Y173A}$ complex.

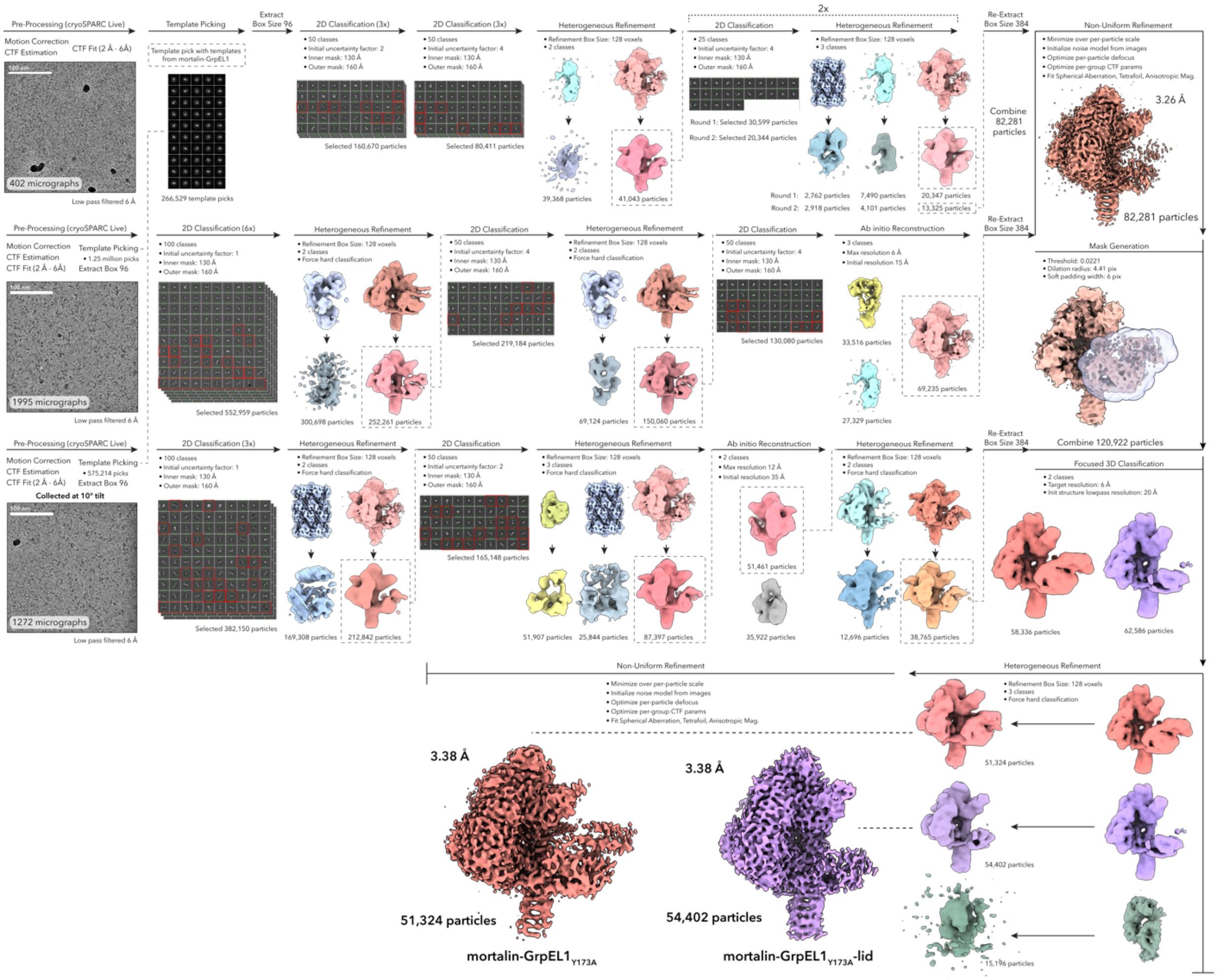


**Supplementary Figure 11. CryoEM data processing of the mortalin-GrpEL1$_{Y173A}$ complexes.** See ***Methods*** for detailed explanation of cryoEM data processing. For 2-D classification classes highlighted in red were excluded from the selected particles. Relevant parameters for each step are listed accordingly.

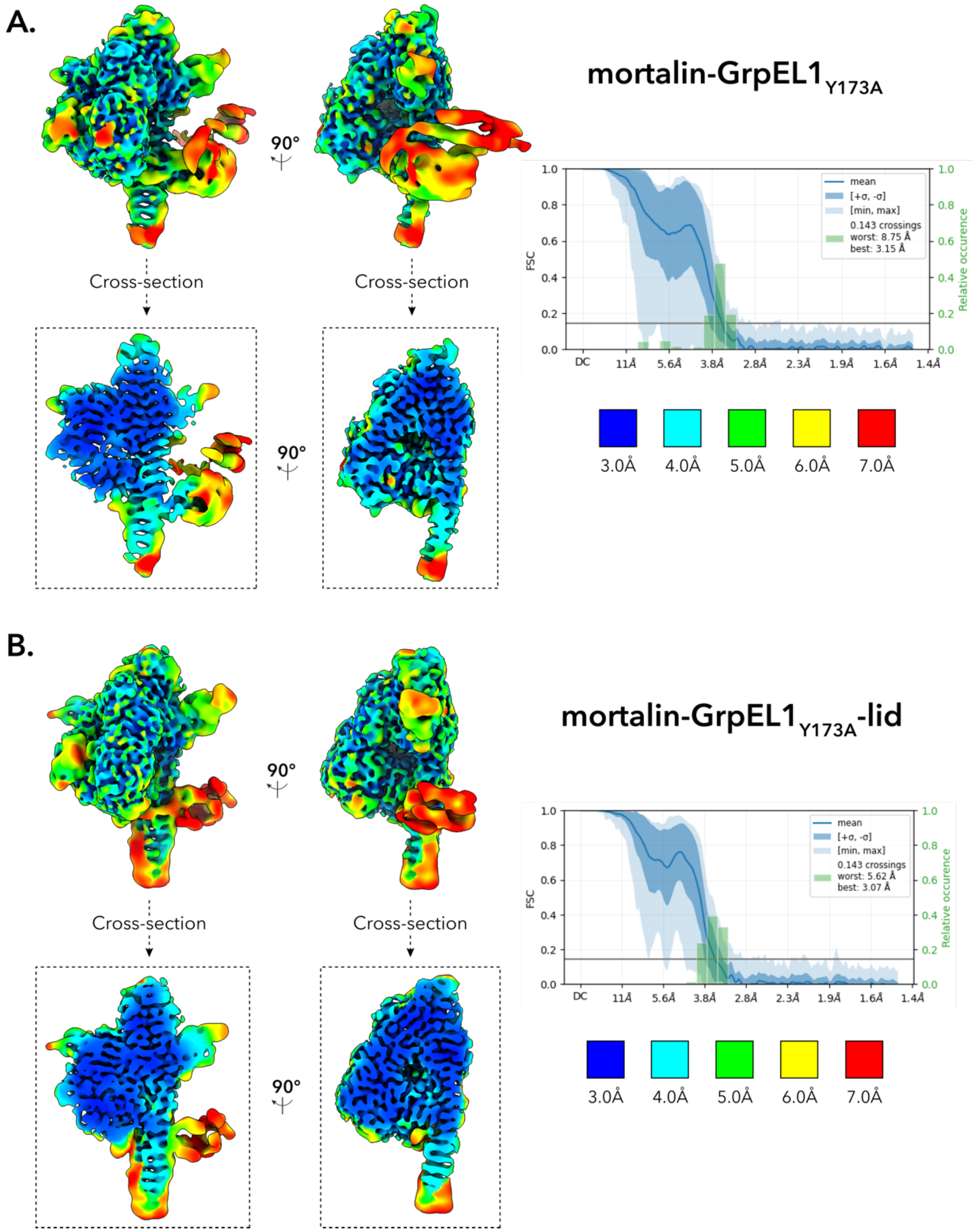


**Supplementary Figure 12. Local resolution estimation of the mortalin-GrpEL1$_{Y173A}$ and mortalin-GrpEL1$_{Y173A}$–lid complexes. A.** Locally filtered EM density of the mortalin-GrpEL1$_{Y173A}$ complex colored by local resolution. Cross-sections of the EM density are shown below each view. 3-D Fourier shell correlation (FSC) plots were generated from the independent half maps contributing to the ~3.38 Å mortalin-GrpEL1$_{Y173A}$ map. **B.** Locally filtered EM density of the mortalin-GrpEL1$_{Y173A}$-lid complex colored by local resolution. Cross-sections of the EM density are shown below each view. 3-D Fourier shell correlation (FSC) plots were generated from the independent half maps contributing to the ~3.38 Å mortalin-GrpEL1$_{Y173A}$-lid map.

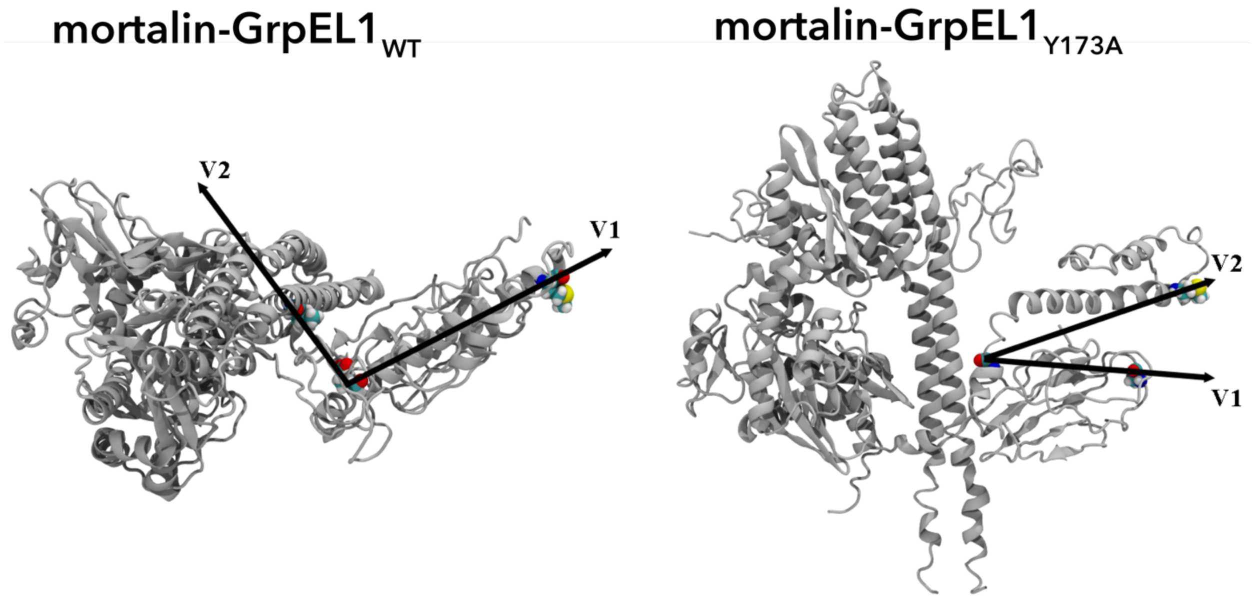


**Supplementary Figure 13. Measurement of the lateral and medial motions of the SBDα lid.**[60] (*left*) Vectors used to measure the lateral motions associated with mortalin-GrpEL1$_{WT}$ in **Figure 7C**. V1 is defined from the Cα of residue 570 to the Cα of residue 596 in mortalin SBDα. V2 is defined from the Cα of residue 570 in SBDα to the Cα of residue 101 in the GrpEL1-B long α-helix. (*right)* Vectors used to measure the medial motions associated with mortalin-GrpEL1$_{Y173A}$ in **Figure 7D.** V1 is defined from the Cα of residue 562 to the Cα of residue 511 in mortalin. V2 is defined from the Cα of residue 562 to the Cα of residue 596 in mortalin.

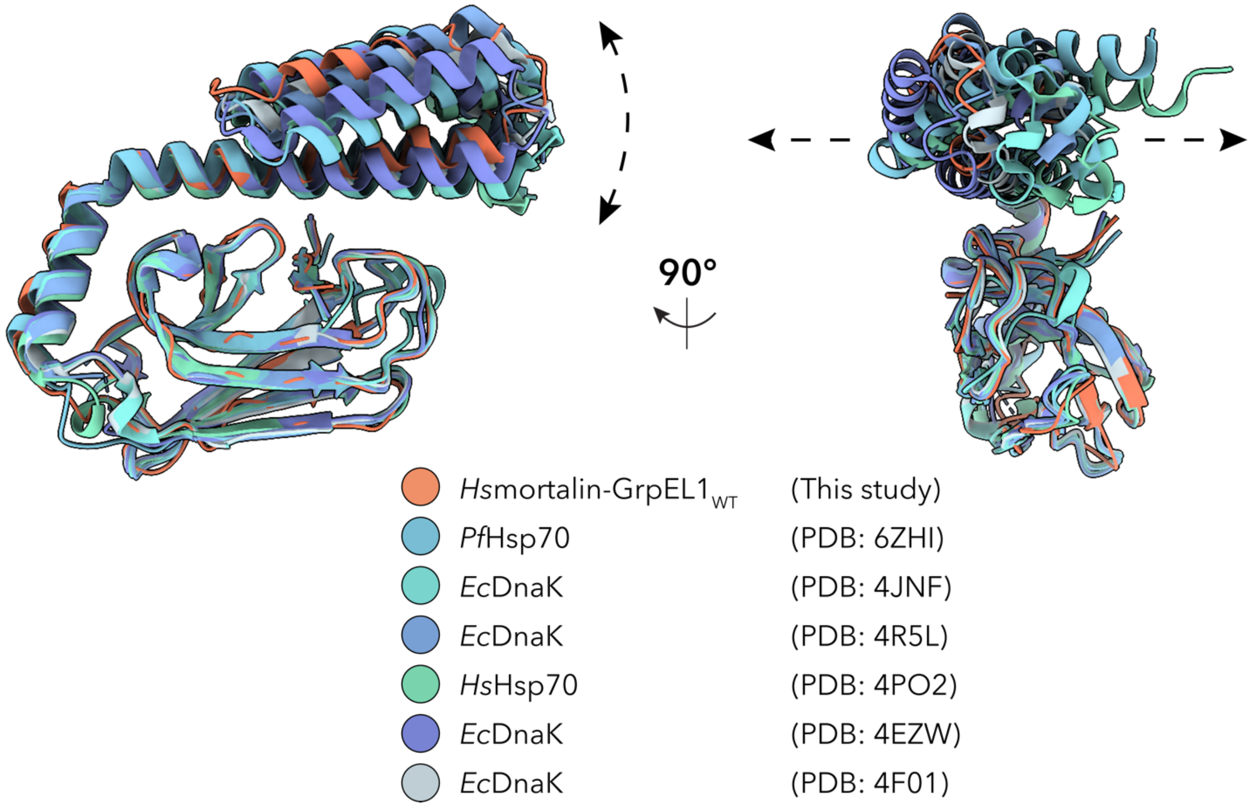


**Supplementary Figure 14. Comparison of SBDα in mortalin-GrpEL1$_{WT}$ with existing Hsp70 crystal structures.** The SBDβ subdomain (residues 440-551) in mortalin-GrpEL1$_{WT}$ were aligned to the SBDβ subdomains of available Hsp70 structures (PDB IDs are indicated in the figure) to exemplify the flexibility in the SBDα across these structures.

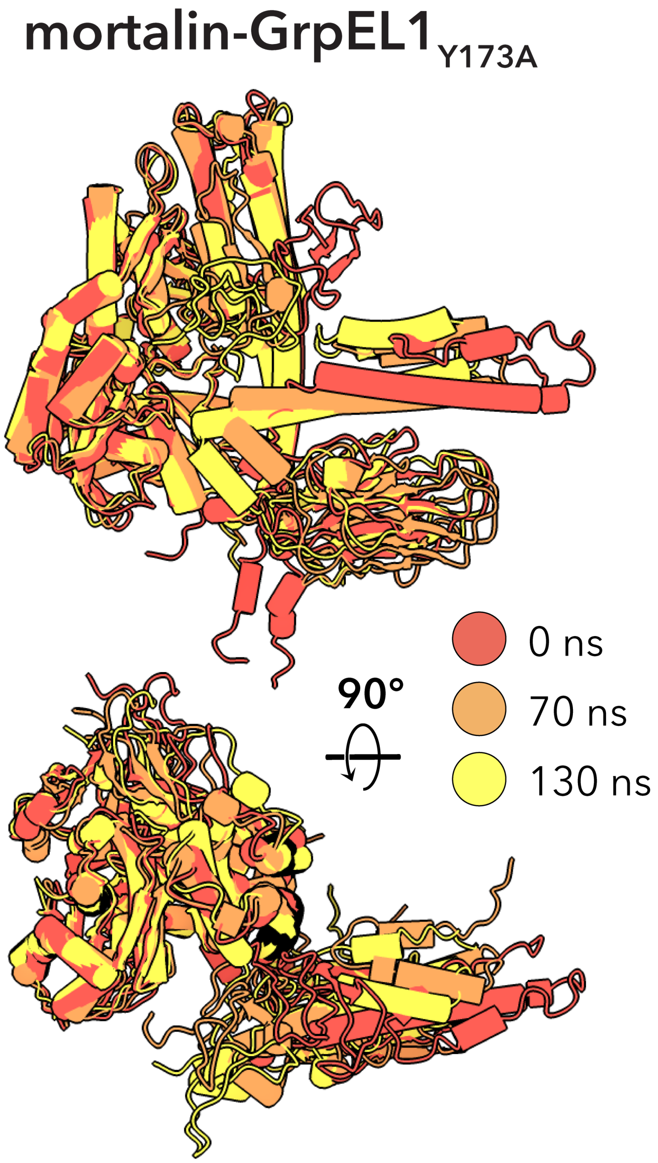


**Supplementary Figure 15. Replicate #3 of the all-atom MD simulations of the mortalin-GrpEL1$_{Y173A}$ complex.** The third replicate of the all-atom MD simulations of mortalin-GrpEL1$_{Y173A}$ described a combination of lateral and medial motions. Timepoints at 0, 70, and 130ns are represented as the start, middle, and end of the simulation.

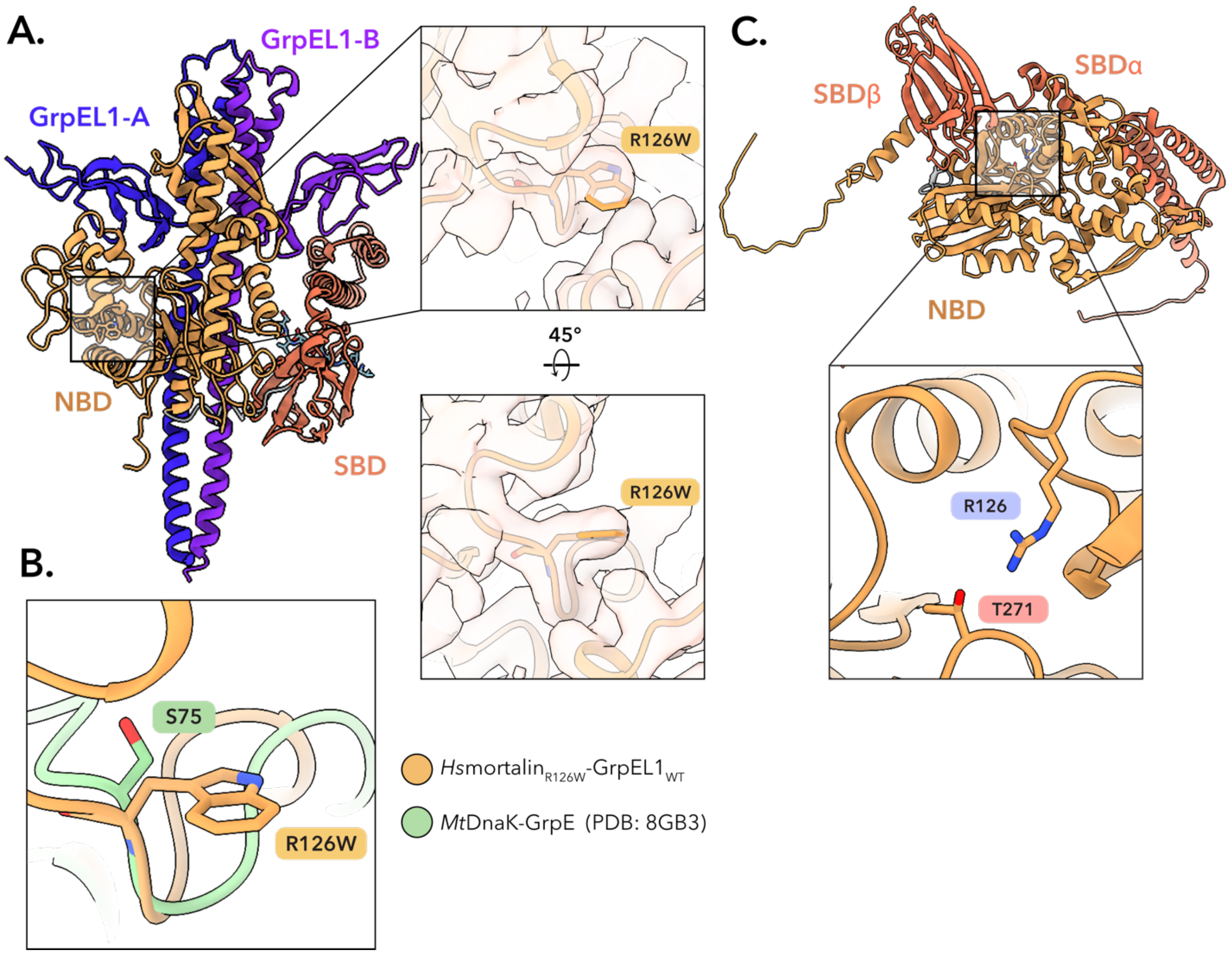


**Supplementary Figure 16. Structural analysis of R126W in the mortalin-GrpEL1$_{WT}$ complex cryoEM structure. A.** R126W does not appear to make additional contacts within the NBD of mortalin when complexed with GrpEL1. **B.** Superposition of *Hs*mortalin-GrpEL1$_{WT}$ with *Mt*DnaK-GrpE (PDB: 8GB3). S75 in *Mt*DnaK, corresponding to R126 in *Hs*mortalin, does not appear to contribute to NBD stabilization or complex formation in *Mt*DnaK-GrpE. **C.** AlphaFold2 predicted structure of *Hs*mortalin. In the apo-nucleotide state, R126 may interact with T271 to stabilize interactions across the IB and IIB NBD lobes that would be absent in the R126W mutant.

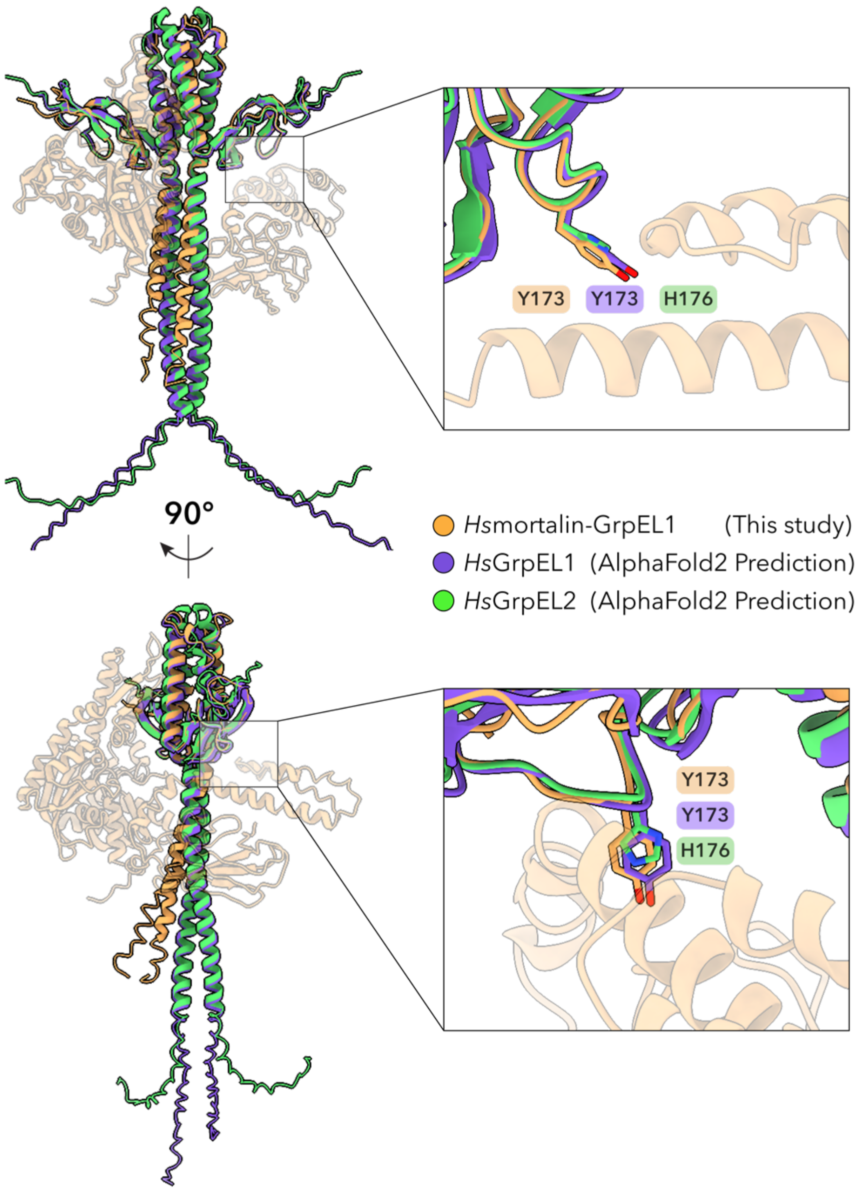


**Supplementary Figure 17. Structural comparisons between mortalin-GrpEL1$_{WT}$, *Hs*GrpEL1, and *Hs*GrpEL2.** GrpEL1-A and GrpEL1-B in the mortalin-GrpEL1$_{WT}$ structure were aligned to the AlphaFold2[47] predicted structures of *Hs*GrpEL1 and *Hs*GrpEL2. While Y173 is substituted for H176 in GrpEL2, the positioning of H176 is analogous to Y173 in the mortalin-GrpEL1$_{WT}$ and GrpEL1 predicted structures.

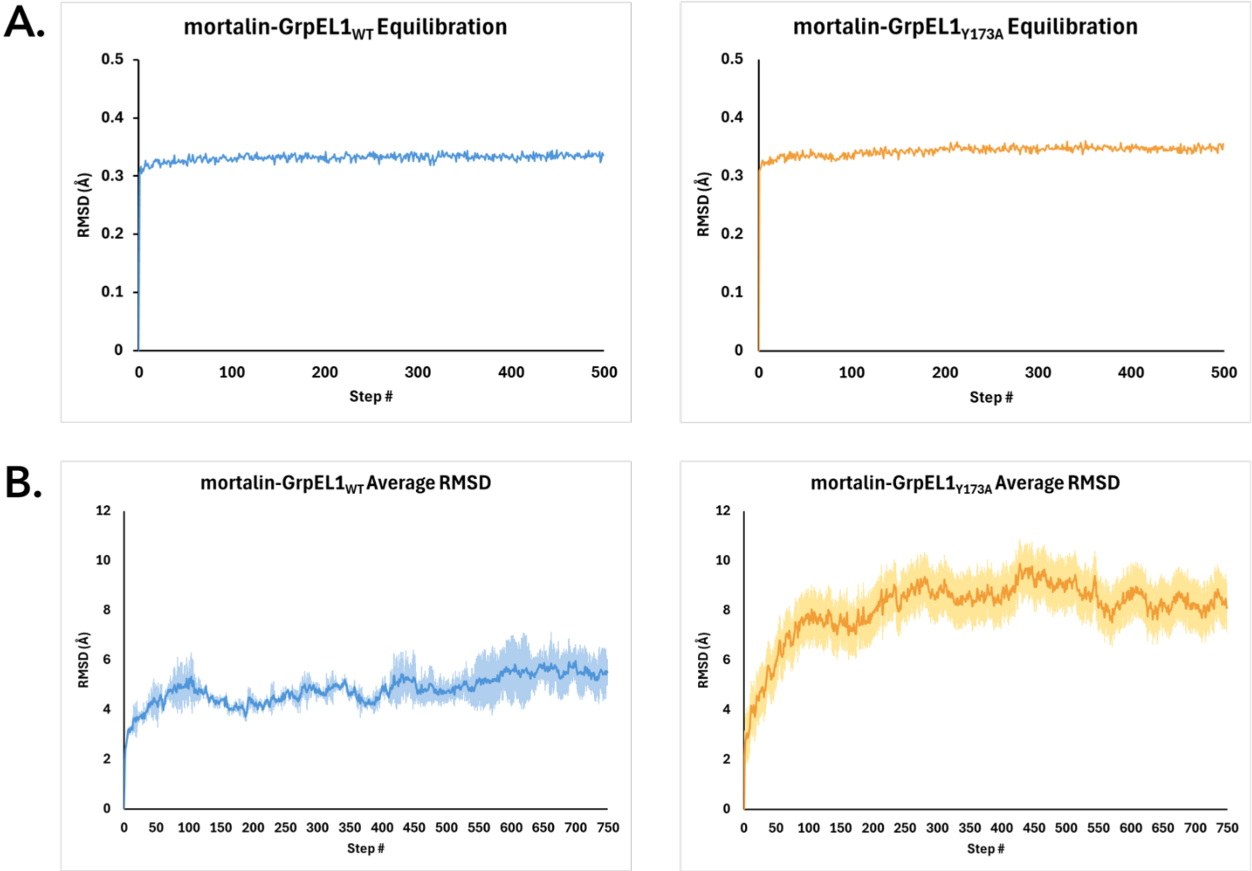


**Supplementary Figure 18. Equilibration and production RMSD analysis of the mortalin-GrpEL1$_{WT}$ and mortalin-GrpEL1$_{Y173A}$ all-atom molecular dynamics simulations.[55] A.** Equilibration for mortalin-GrpEL1$_{WT}$ (*left)* and mortalin-GrpEL1$_{Y173A}$ *(right)* prior to production randomization in triplicate. **B.** Production RMSD analysis of mortalin-GrpEL1$_{WT}$ (*left)* and mortalin-GrpEL1$_{Y173A}$ (*right)* throughout the 150ns simulation. Shaded regions represent the standard deviation across the triplicates.

**Supplementary Movie 1: Anisotropic network modelling (ANM) analysis of mortalin-GrpEL1$_{WT}$.** Ten ANM modes of the mortalin-GrpEL1$_{WT}$ structure were visualized using VMD[58] and highlight the flexibility of the mortalin SBD. Areas of high motion are colored in red, and areas of low motion are colored in blue.

**Supplementary Movie 2: Anisotropic network modelling (ANM) analysis of mortalin-GrpEL1$_{Y173A}$.** Ten ANM modes of the mortalin-GrpEL1$_{Y173A}$ structure were visualized using VMD[58] and highlight the flexibility of the mortalin SBD. Areas of high motion are colored in red, and areas of low motion are colored in blue.

| | Mortalin-GrpEL1$_{WT}$ | Mortalin-GrpEL1$_{Y173A}$ | |
|---|---|---|---|
| **Data Collection** | | | |
| Magnification | 130kx | 165kx | |
| Voltage (kV) | 300 | 300 | |
| Spherical Aberration (mm) | 2.7 | 2.7 | |
| Electron Exposure ($e^-/Å^2$) | 60 | 60 | |
| Defocus range ($\mu$m) | -1.0 to -2.5 | -1.0 to -2.5 | |
| Pixel size (Å, Physical/Digital) | 0.935 | 0.735 | |
| Energy Filter Slit Width (eV) | 10 | 10 | |
| Movies | 4570 | 3669 | |
| **Map Statistics and Post-Processing** | | **Mortalin-GrpEL1$_{Y173A}$** | **Mortalin-GrpEL1$_{Y173A}$-lid** |
| Accession Codes (EMDB, PDB) | EMD-44675, 9BLS | EMD-44676, 9BLT | EMD-44677, 9BLU |
| Symmetry imposed | C1 | C1 | C1 |
| Map Resolution (Å) | 2.96 | 3.38 | 3.38 |
| Local resolution range for 75% of voxels | 5.825 | 7.874 | 7.326 |
| Local resolution range (model) | 2.579 - 31.802 | 3.014 - 52.055 | 2.999 - 47.490 |
| Map sharpening B factor ($Å^2$) | 97.9 | 83.2 | 85.7 |
| Map sharpening method | DeepEMhancer | DeepEMhancer | DeepEMhancer |
| Table Name: | table1 | table1 | table1 |
| **Model Statistics and Validation** | | | |
| Model composition | | | |
| Non-hydrogen atoms | 7113 | 7099 | 6378 |
| Protein | 921 | 921 | 831 |
| Nucleic acids | 0 | 0 | 0 |
| Ligands | 0 | 0 | 0 |
| Waters | 0 | 0 | 0 |
| R.M.S deviations | | | |
| Length (Å) | 0.003 | 0.003 | 0.003 |
| Angles (°) | 0.503 | 0.505 | 0.452 |
| MolProbity score | 2.41 | 2.11 | 1.92 |
| MolProbity Clashscore | 8.55 | 8.29 | 6.89 |
| CaBLAM (% outliers) | 2.32 | 1.66 | 2.21 |
| Rotamer outliers (%) | 5.30 | 2.72 | 3.18 |
| Cis peptides (#, %) | 0.0/0.0 | 0.0/0.0 | 0.0/0.0 |
| Ramachandran Plot | | | |
| Favored | 94.09 | 95.18 | 97.08 |
| Allowed | 5.91 | 4.82 | 2.92 |
| Outliers | 0.0 | 0.0 | 0.0 |

**Table 1. CryoEM data collection and refinement statistics of mortalin-GrpEL1$_{WT}$, mortalin-GrpEL1$_{Y173A}$, and mortalin-GrpEL1$_{Y173A}$-lid.**
